# HUWE1 controls tristetraprolin proteasomal degradation by regulating its phosphorylation

**DOI:** 10.1101/2022.08.29.505645

**Authors:** Sara Scinicariello, Adrian Söderholm, Markus Schäfer, Alexandra Shulkina, Irene Schwartz, Kathrin Hacker, Rebeca Gogova, Robert Wolfgang Kalis, Kimon Froussios, Valentina Budroni, Annika Bestehorn, Tim Clausen, Pavel Kovarik, Johannes Zuber, Gijs A. Versteeg

**Affiliations:** Department of Microbiology, Immunobiology and Genetics, Max Perutz Labs, University of Vienna, Vienna BioCenter (VBC), Vienna, Austria; Vienna BioCenter PhD Program, Doctoral School of the University of Vienna and Medical University of Vienna, Vienna BioCenter (VBC), Vienna, Austria; Research Institute of Molecular Pathology (IMP), Vienna BioCenter (VBC), Vienna, Austria; Medical University of Vienna, Vienna BioCenter (VBC), Vienna, Austria

## Abstract

Tristetraprolin (TTP) is a critical negative immune regulator. It binds AU-rich elements in the untranslated-regions of many mRNAs encoding pro-inflammatory mediators, thereby accelerating their decay. A key but poorly understood mechanism of TTP regulation is its timely proteolytic removal: TTP is degraded by the proteasome through yet unidentified phosphorylation-controlled drivers. In this study, we set out to identify factors controlling TTP stability. Cellular assays showed that TTP is strongly lysine-ubiquitinated, which is required for its turnover. A genetic screen identified the ubiquitin E3 ligase HUWE1 as a strong regulator of TTP proteasomal degradation, which we found to control TTP stability indirectly by regulating its phosphorylation. Pharmacological assessment of multiple kinases revealed that HUWE1-regulated TTP phosphorylation and stability was independent of the previously characterized effects of MAPK-mediated S52/S178 phosphorylation. HUWE1 function was dependent on phosphatase and E3 ligase binding sites identified in the TTP C-terminus. Our findings indicate that while phosphorylation of S52/S178 is critical for TTP stabilization at earlier times after pro-inflammatory stimulation, phosphorylation of the TTP C-terminus controls its stability at later stages.

## Introduction

Dynamic regulation of the immune system is essential to mount a defense against pathogens upon infection, yet shut-off the response at the appropriate time during resolution. Since most cytokines and other pro-inflammatory mediators are transcriptionally induced during infection, an essential aspect of returning to homeostatic conditions is the timely removal of their mRNAs during resolution.

Tristetraprolin (TTP; also known as ZFP36 or TIS11A) is an RNA-binding protein that interacts with AU-rich elements (ARE) present in the 3’-untranslated-regions (UTR) of many mRNAs encoding pro-inflammatory mediators (Sedlyarov *et al*, 2016; Galloway *et al*, 2016; Lai *et al*, 1999). Subsequently, TTP recruits the CCR4-NOT decapping and deadenylation complex to target mRNAs, resulting in their destabilization and removal from the cell (Sandler *et al*, 2011; Lykke-Andersen & Wagner, 2005; Fabian *et al*, 2013; Tiedje *et al*, 2016; Lai *et al*, 2003). TTP binds to the AREs of a multitude of mRNAs encoding cytokines and other immune-related factors, yet not all of them are destabilized (Sedlyarov *et al*, 2016; Tiedje *et al*, 2016; Zhang *et al*, 2017; Moore *et al*, 2018). This has suggested that additional - hitherto unknown-regulatory mechanisms are at play controlling TTP-dependent mRNA degradation, which may differ in various cell types.

The biological importance of TTP for proper dampening of the inflammatory response is underpinned by the observation that *Zfp36* (the gene encoding TTP)-deficient mice develop systemic inflammation characterized by arthritis, dermatitis, conjunctivitis and cachexia, which has been coined TTP deficiency syndrome (Taylor *et al*, 1996). One of the main deregulated ARE-containing mRNAs driving the inflammatory phenotype in *Zfp36*-deficient mice is *Tnf* (Taylor *et al*, 1996; Carballo *et al*, 1998), although additionally *Il1a/b, Il23*, and *Ccl3* have been implicated as well (Sneezum *et al*, 2020; Molle *et al*, 2013; Kang *et al*, 2011).

TTP itself is regulated at the transcriptional, post-transcriptional, and post-translational levels. Most cell types express low levels of *Zfp36* mRNA in unstimulated conditions, the transcription of which is robustly induced by proinflammatory stimuli including the Toll-like receptor 4 (TLR4) agonist lipopolysaccharide (LPS) in myeloid cells such as macrophages (Carballo *et al*, 1998; Suzuki *et al*, 2003; Sauer, 2006; Lai *et al*, 1995; Schaljo *et al*, 2009). At the post-translational level, TTP is phosphorylated at over 30 residues by inflammation-activated stress kinases (Hitti *et al*, 2006; Brook *et al*, 2006; Clark & Dean, 2016; Ronkina *et al*, 2019).

The biological relevance of most TTP phospho-sites and the identity of the involved kinases remain unknown (Clark & Dean, 2016; Ronkina *et al*, 2019). Most characterized are phosphorylation events at residues S52 and S178 in murine TTP that are mediated by the inflammation-activated kinase MK2, which acts down-stream of p38 mitogen-activated protein kinase (MAPK) (Hitti *et al*, 2006; Brook *et al*, 2006). In mice, TTP mutants lacking these phosphorylation sites are highly unstable and rapidly degraded, yet highly biologically active (Ross *et al*, 2015).

This has given rise to a model in which TTP is predominantly unphosphorylated and rapidly degraded in unstimulated cells, whereas pro-inflammatory cell signaling not only increases *Zfp36* transcription, but also TTP S52/S178 phosphorylation and stabilization through interaction with 14-3-3 proteins (Sedlyarov *et al*, 2016; Kratochvill *et al*, 2011). However, in this S52/S178 phosphorylated state TTP is thought to be inactive, whereas during dephosphorylation of these residues at later times in the inflammatory response, TTP actively mediates mRNA degradation (Sedlyarov *et al*, 2016; Kratochvill *et al*, 2011). Nevertheless, the impact of the other 30+ phosphorylated residues on TTP stability and activity has remained largely elusive.

Proteasomes are the main degradation machines of cells for homeostatic protein turn-over (Bard *et al*, 2018). 20S core particles contain catalytic activity, yet lack receptors for ubiquitin and ATPse activity for unfolding and translocation of proteins into the catalytic chamber. Association of 19S regulatory particles containing ubiquitin receptors and AAA+ ATPase activity assembles 26S proteasomes, which are considered the main degradative entities for poly-ubiquitinated proteins (Bard *et al*, 2018).

TTP protein is degraded through the proteasome, as previous studies showed that 20S proteasome inhibition stabilizes TTP. Moreover, a previous study suggested that TTP may be directly degraded by 20S proteasomes in a ubiquitin-independent manner. In this context, important destabilizing intrinsically-disordered regions in the N and C termini of TTP were identified and have been suggested to putatively allow direct degradation by 20S proteasomes (Brook *et al*, 2006; Ngoc *et al*, 2014). Yet, other regulators of intra-cellular TTP protein abundance have remained elusive. In this study, we set out to identify and characterize novel factors that control TTP turn-over, thereby affecting pro-inflammatory output.

Through genetic loss-of-function screening, we identified several novel determinants of TTP abundance, including the giant ubiquitin E3 ligase HUWE1. Our data indicate that TTP is strongly poly-ubiquitinated on lysines in its zinc finger domain, and degraded by the proteasome in a ubiquitin-dependent manner. Moreover, we identified a novel role for the E3 ligase HUWE1 in indirectly controlling TTP turn-over through mediating its phosphorylation via multiple stress kinases, and reduced dephosphorylation.

## Results

### TTP is degraded in a ubiquitin-dependent manner

Pro-inflammatory stimuli such as LPS drive both transcription of *Zfp36* (the gene encoding TTP), and phosphorylation of the TTP protein. To study how TTP protein levels are regulated, we established a macrophage cell line expressing exogenous TTP from a constitutively active promoter, uncoupling *Zfp36* transcription from regulatory effects on TTP protein stability in the absence or presence of pro-inflammatory signals.

Consistent with previous studies, endogenous TTP protein was rapidly induced by LPS (Carballo *et al*, 1998; Suzuki *et al*, 2003; Sauer, 2006; Lai *et al*, 1995; Schaljo *et al*, 2009), and in the absence of *de novo* protein synthesis, rapidly degraded (Fig. 1A) (Brook *et al*, 2006; Ngoc *et al*, 2014). Treatment with proteasome inhibitor MG132 almost completely prevented TTP degradation, indicating that its degradation is predominantly through the proteasome. Under these conditions a hyper-phosphorylated form of TTP accumulated, suggesting that phosphorylation is important for regulation of its stability.

**Figure 1.**
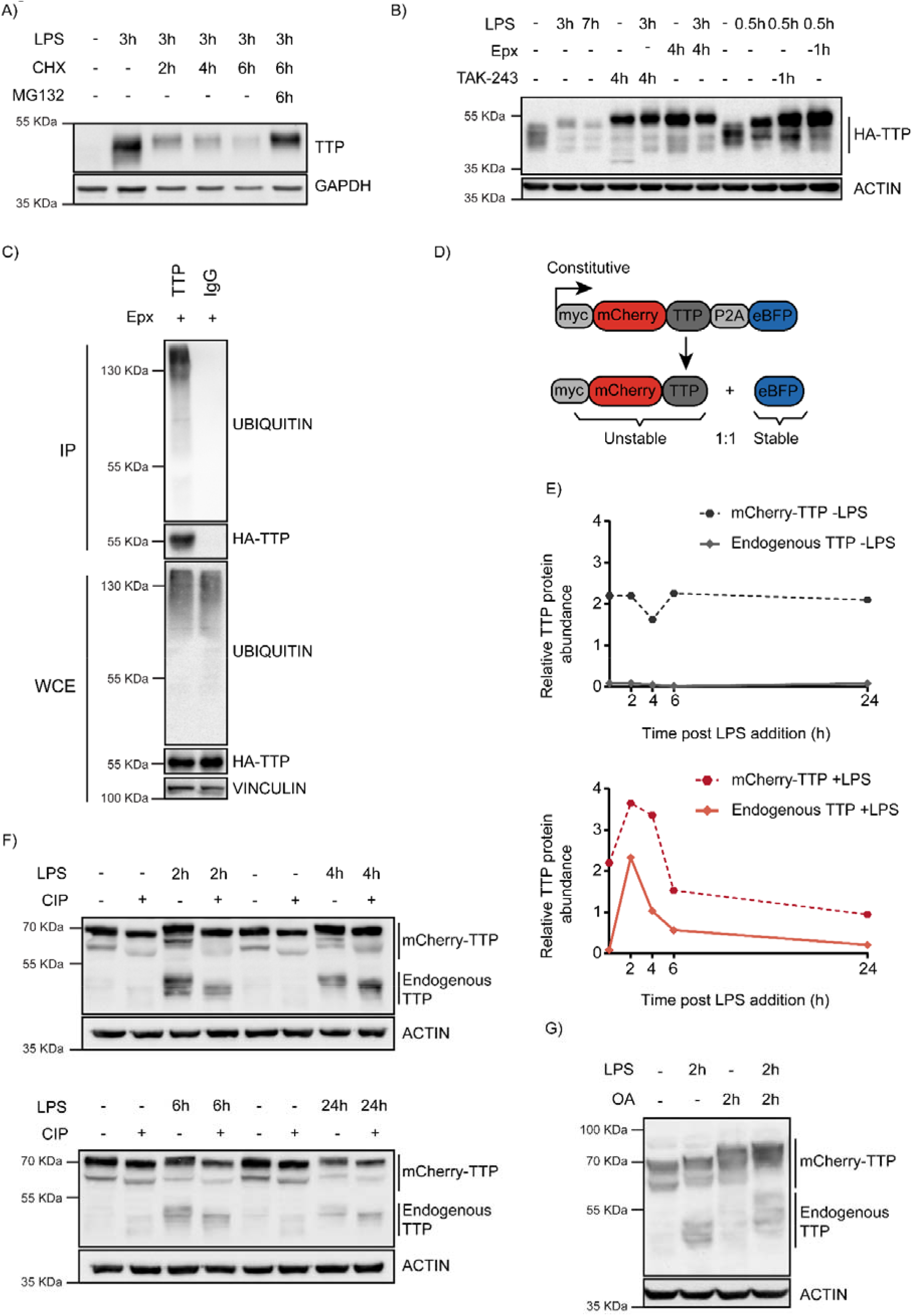
TTP is degraded by the proteasome in a ubiquitin-dependent manner. **A)** RAW264.7 murine macrophages were stimulated with LPS and incubated with the translation inhibitor cycloheximide (CHX) and the proteasome inhibitor MG132 for the indicated times (h), after which TTP levels were analyzed by western blot. **B)** 3xHA-TTP-expressing RAW264.7 cells were incubated with LPS or left unstimulated. Cells were then treated with E1 enzyme inhibitor (TAK-234) or the proteasome inhibitor Epoxomicin (Epx). Protein levels were assessed by western blot. **C)** RAW264.7 cells stably expressing 3xHA-tagged TTP protein were treated with Epx for 5 h, after which TTP was immunoprecipitated, and its ubiquination analysed by western blot. **D)** Schematic representation of the TTP stability reporter construct. Constitutively expressed myc-tagged mCherry-TTP fusion protein and enhanced blue fluorescent protein (eBFP2) are translated at equimolar levels through a P2A site. **E-F)** RAW264.7-Dox-Cas9-mCherry-TTP cells were stimulated with LPS for the indicated times. Subsequently, cell lysates were treated with Calf Intestinal Phosphatase (CIP) for 2 h. at 37 °C, and TTP levels analyzed by western blot. Non-saturated western blot signals for mCherry-TTP and endogenous TTP protein were quantified, normalized to ACTIN levels, and plotted. **G)** RAW264.7-Dox-Cas9-mCherry-TTP cells where treated with LPS for 2 h, after which PP1/2 inhibitor Okadaic Acid (OA) was added to the culture medium for 2 h. TTP electrophoretic mobility was assessed by western blot.

To investigate whether pro-inflammatory stimuli are exclusively stabilizing TTP, or also provide degradation signals, a macrophage cell line stably expressing HA-tagged TTP was established (Fig. 1B). Under non-stimulated conditions, HA-TTP was detected as medium-range MW species migrating at and above its predicted MW of 36.7 kDa, consistent with it being partially phosphorylated under non-stimulated conditions. Stimulation with LPS resulted in rapid TTP stabilization after 30 min., followed by a reduction of its protein levels at 3 h. and 7 h. post-treatment (Fig. 1B; lanes 2-3). This suggested that pro-inflammatory stimuli may also provide the signaling required for TTP turn-over at longer stimulation times, possibly through regulating its phosphorylation.

To determine whether TTP proteasomal degradation was mediated by ubiquitination, cells were treated with the ubiquitin E1 inhibitor TAK-243, which inhibits *de novo* ubiquitination (Fig. 1B and EV1A). This stabilized TTP under baseline and LPS-stimulated conditions (Fig. 1B), demonstrating that TTP is degraded in a ubiquitination-dependent manner. Consistent with this notion, TTP in denaturing lysates from these cells was detected to be ubiquitinated (Fig. 1C). Moreover, a TTP mutant in which all of its five lysine residues in the TTP zinc finger domain were mutated to arginines (KtoR), accumulated at higher steady-state levels and was substantially less ubiquitinated (Fig. EV1B). Together, these data indicate that TTP is covalently poly-ubiquitinated on one or more lysine residues in its zinc finger domain, marking it for proteasomal degradation.

To enable identification of TTP abundance regulators by genetic screening, a macrophage cell line stably expressing unstable mCherry-TTP and stable BFP was established (Fig. 1D). The stable BFP served as an internal control, as it is translated in equimolar amounts from the same transcript through a P2A ribosomal skip site. mCherry-TTP accumulated in cells as a stable protein under non-stimulated conditions (Fig. 1E; top panel). In contrast, LPS stimulation initially further stabilized mCherry-TTP, yet subsequently facilitated its degradation, phenocopying its endogenous TTP counterpart (Fig. 1E; bottom panel, and EV1C). Treatment of lysates from these cells with alkaline phosphatase collapsed higher migrating endogenous and exogenous TTP species (Fig. 1F), whereas inhibition of the phosphatases PP1 and PP2 by okadaic acid (OA) increased them (Fig. 1G), indicating that mCherry-TTP is phosphorylated in a similar fashion as endogenous TTP.

Together, these data show that LPS-stimulation initially stabilizes TTP, whereas at later time points its induced cell signaling events direct TTP degradation.

### The E3 ligase HUWE1 is a major determinant of cellular TTP protein abundance

Next, we set out to identify cellular factors regulating TTP protein abundance. To this end, a RAW264.7 mouse macrophage cell line with Dox-inducible Cas9 was established, which in addition expresses mCherry-TTP (Fig. 1D and 2A). To enable identification of essential genes, a cell line was established which only functionally edits in the presence of doxycycline (Dox), but not in its absence (Appendix 1A).

A genome-wide lentiviral sgRNA library was transduced into these cells, ensuring one integration per cell. Knock-outs were induced by treatment with Dox for three and six days to identify regulators irrespective of essential gene functions and different protein half-lives. Subsequently, TTP was destabilized by LPS treatment, cells with high and low mCherry-TTP content were sorted, and their integrated sgRNA coding sequences quantified by next generation sequencing (Fig. 2A, and Appendix Fig. S2). In parallel, sorted cells with high or low levels of the stable BFP control (Fig. 1D, and Appendix Fig. S2) were likewise processed, and used for identifying non-specific factors.

**Figure 2.**
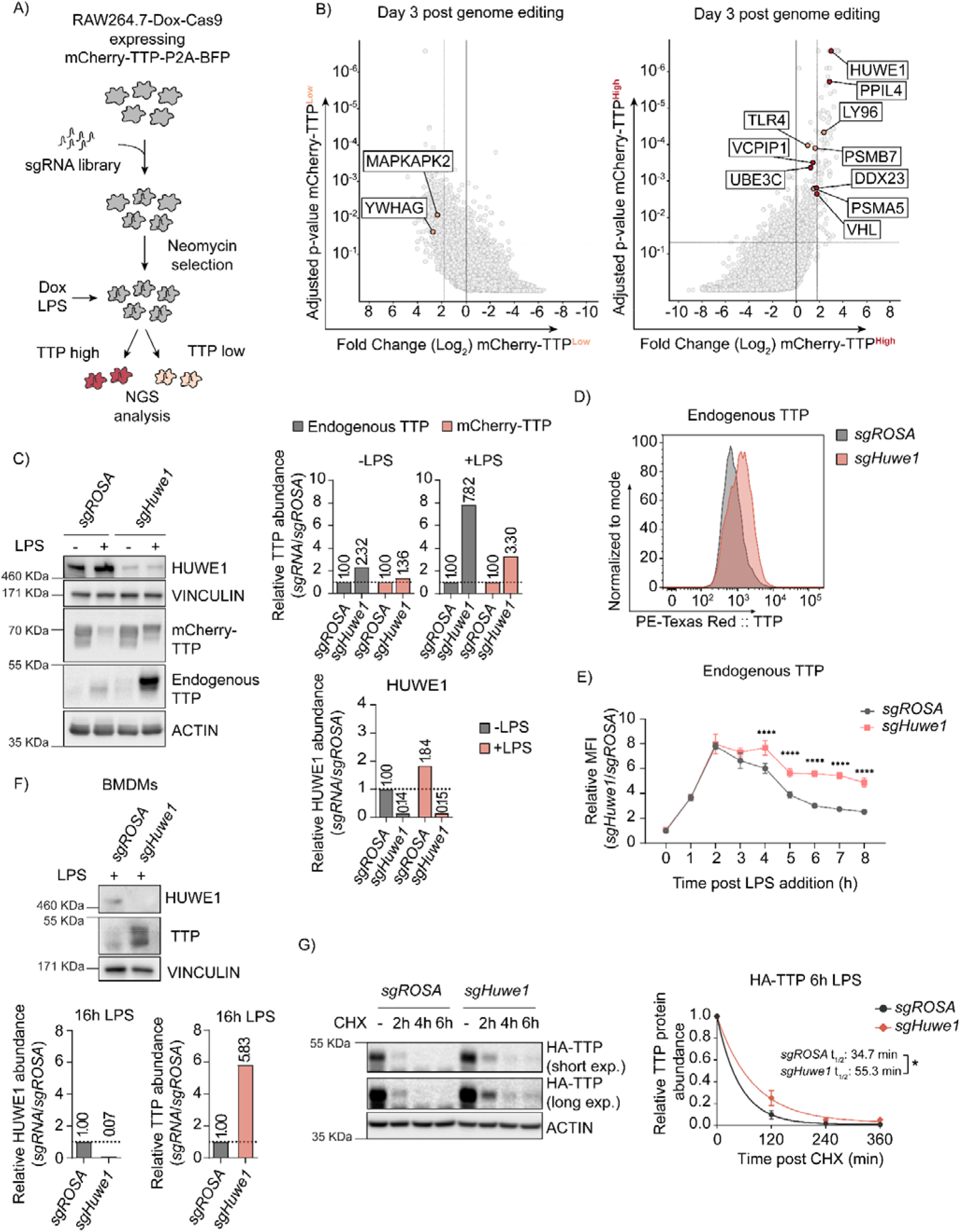
Genome-wide CRISPR-Cas9 knockout screen identified the E3 ligase HUWE1 as a regulator of TTP stability. **A)** Overview of FACS-based CRISPR-Cas9 knockout screening procedure using the RAW264.7-Dox-Cas9-mCherry-TTP cell line. Cells expressing high and low levels of mCherry-TTP protein were sorted, and their integrated sgRNA coding sequences determined by next generation sequencing. **B)** Read counts per million in the mCherry-TTP^high^ cells at 3 days after Cas9 induction were compared to those in unsorted cells from the same day, sgRNA enrichment calculated by MAGeCK analysis, and log_2_-fold change and adjusted p-value plotted. Genes enriched in the sorted populations that met the following criteria are indicated in red: a log_2_ fold-change of < 1.8 (mCherry-TTP^low^) or > 1.8 (mCherry-TTP^high^), adjusted p-value < 0.05, not enriched in the matching eBFP2^low^ or eBFP2^high^ sorted cells. **C)** Cas9 was induced with Dox for 5 days in RAW264.7-Dox-Cas9 cells expressing either sg*ROSA* or sg*Huwe1*. Subsequently, cells were treated with LPS for 16 h, and TTP protein levels were assessed by western blot. HUWE1, mCherry-TTP and endogenous TTP abundance was quantified and plotted. The TTP and ACTIN panels are the left four lanes from the blot presented in Fig. EV2B. **D)** RAW264.7-Dox-Cas9 cells expressing sg*ROSA* or sg*Huwe1* were treated with Dox for 5 days to induce Cas9. Then, cells were incubated with LPS for 16 h. or left unstimulated, and endogenous TTP protein levels analyzed by intracellular staining, followed by flow cytometry. **E)** sg*ROSA-* or sg*Huwe1-*targeted RAW264.7-Dox-Cas9 cells were treated with LPS for the indicated times (h) and TTP levels were analyzed by flow cytometry. Normalized mean fluorescent intensity (MFI) was plotted. Data represent the mean and s.d.; n = 3. \*\*\*\**p* ≤ 0.0001. **F)** Bone marrow-derived macrophages (BMDMs) isolated from Cas9-expressing knock-in mice were stably transduced with sg*ROSA* or sg*Huwe1*. Cells were incubated with LPS for 16 h. or left unstimulated. Endogenous TTP protein levels were determined by western blot. Quantified TTP levels normalized to VINCULIN are plotted. **G)** sg*ROSA-* or sg*Huwe1-*RAW264.7-Dox-Cas9 cells stably expressing 3xHA-TTP were treated for 6 h. with LPS, followed by CHX. Protein lysates were harvested at the indicated time points (h). TTP levels were measured by western blot, quantified, plotted, and TTP half-life calculated. n = 2, *p ≤ 0.05.

As anticipated, factors previously reported to be important for stabilizing TTP (e.g. *Mapkapk2*/*Mk2, Ywhag*/*14-3-3γ, Mapk14*/*p38*) were significantly enriched in the mCherry-TTP^low^ cell pool (Table EV1, Fig. 2B and EV2A; left and top panels, respectively). Consistent with mCherry-TTP proteasomal degradation being LPS-dependent (Fig. 1E), key factors for TLR4-signaling (*Tlr4, Ly96/Md2*), and components of the proteasome (*Psma5, Psmb7*) were significantly enriched in mCherry-TTP^high^ sorted cells (Fig. 2B and EV2A; right and bottom panels, respectively). Moreover, various additional new candidates controlling cellular TTP abundance were identified, including the giant ubiquitin E3 ligase *Huwe1* (Fig. 2B and EV2A, right and top panels, respectively). Individual targeting of these candidates increased endogenous TTP and exogenous mCherry-TTP protein levels by western blot (Fig. EV2B), attesting to the validity and predictive quality of our screen.

In particular, HUWE1 was identified as a strong determinant of endogenous and exogenous TTP protein abundance by western blot (Fig. 2C and EV2B; compare LPS-treated samples), and flow cytometry (Fig. 2D), without affecting *Zfp36* mRNA levels (Fig. EV2C). Consistent with an increase in protein stability, inducible *Huwe1* knock-out significantly increased endogenous TTP protein levels at later time points post-LPS stimulation (Fig. 2E). Given that after initial stabilization, LPS mediates TTP degradation (Fig. 1E), we tested whether *Huwe1* knock-out affected TLR4 signaling. To this end, the effect of *Huwe1* loss on IκBα, which is degraded in a proteasome-dependent manner upon LPS stimulation (Fig. EV2D), was measured. *Huwe1* knock-out increased TTP protein levels (Fig. EV2E; top panel), but did neither affect LPS-induced degradation of mCherry-IκBα by flow cytometry (Fig. EV2E; bottom panel), nor endogenous IκBα by western blot (Fig. EV2F). This shows that the *Huwe1* knock-out does not affect cell signaling between TLR4 and IκBα, and this does not contribute to TTP stabilization in *Huwe1*-deficient cells.

Next, we determined whether the HUWE1-dependent control of TTP abundance in the RAW264.7 mouse macrophage cell line was conserved across species and cell types. To this end, *HUWE1* was targeted in the human colon carcinoma cell line RKO, which - unlike most myeloid cells-have low detectable levels of TTP in the absence of any stimulation (Fig. EV2G). Similar to the phenotype in RAW264.7 cells, *HUWE1* knock-out in RKO cells strongly increased TTP protein levels (Fig. EV2G), indicating that HUWE1 has a similar role in human, non-myeloid cells independent of the TLR4 axis. Moreover, targeting of *Huwe1* in mouse bone marrow derived macrophages (BMDMs), likewise strongly increased high and low molecular weight species of endogenous TTP (Fig. 2F), indicating that the biological importance of *Huwe1* for TTP abundance is relevant in primary cells.

Lastly, we measured whether *Huwe1* ablation affected TTP protein half-life. To this end, control-targeted *sgROSA* and *sgHuwe1* cells were chased in the presence of translation inhibitor cycloheximide (CHX), TTP protein levels analyzed by western blot, and single-step exponential decay curves plotted. *Huwe1* knock-out increased TTP protein half-life by 83% from 35 min. to 55 min. (Fig. 2G), demonstrating that loss of *Huwe1* increases TTP protein half-life. Collectively, these data position HUWE1 as a strong, conserved regulator of TTP protein stability.

### Loss of *Huwe1* decreases the half-life of pro-inflammatory mRNAs controlled by TTP

TTP is essential for the degradation of transcripts with AU-rich elements in their 3’-UTR, encoding pro-inflammatory cytokines such as TNF and IL6. Phosphorylation of S52 and S178 stabilizes TTP, yet reduces its degradation of mRNAs. In contrast, the effect of phosphorylation on other sites has remained elusive. Therefore, we reasoned that increased TTP protein levels upon *Huwe1* ablation could **i)** either result in increased intracellular TTP protein concentrations, and consequently diminished levels of transcripts encoding pro-inflammatory cytokines, or -as a consequence of increased TTP phosphorylation- **ii)** decrease the bio-active pool of TTP, resulting in equal or increased mRNA levels in *Huwe1* knock-out cells.

To investigate whether increased TTP levels upon *Huwe1* loss are biologically relevant, we measured *Tnf* and *Il6* mRNA levels in *Huwe1*-targeted BMDMs and RAW264.7 cells. Consistent with the fact that non-stimulated cells have very low levels of TTP, *Huwe1* knock-out did not alter baseline *Tnf* (Fig. EV3A), or *Il6* (Fig. 3A/B) mRNA levels.

**Figure 3.**
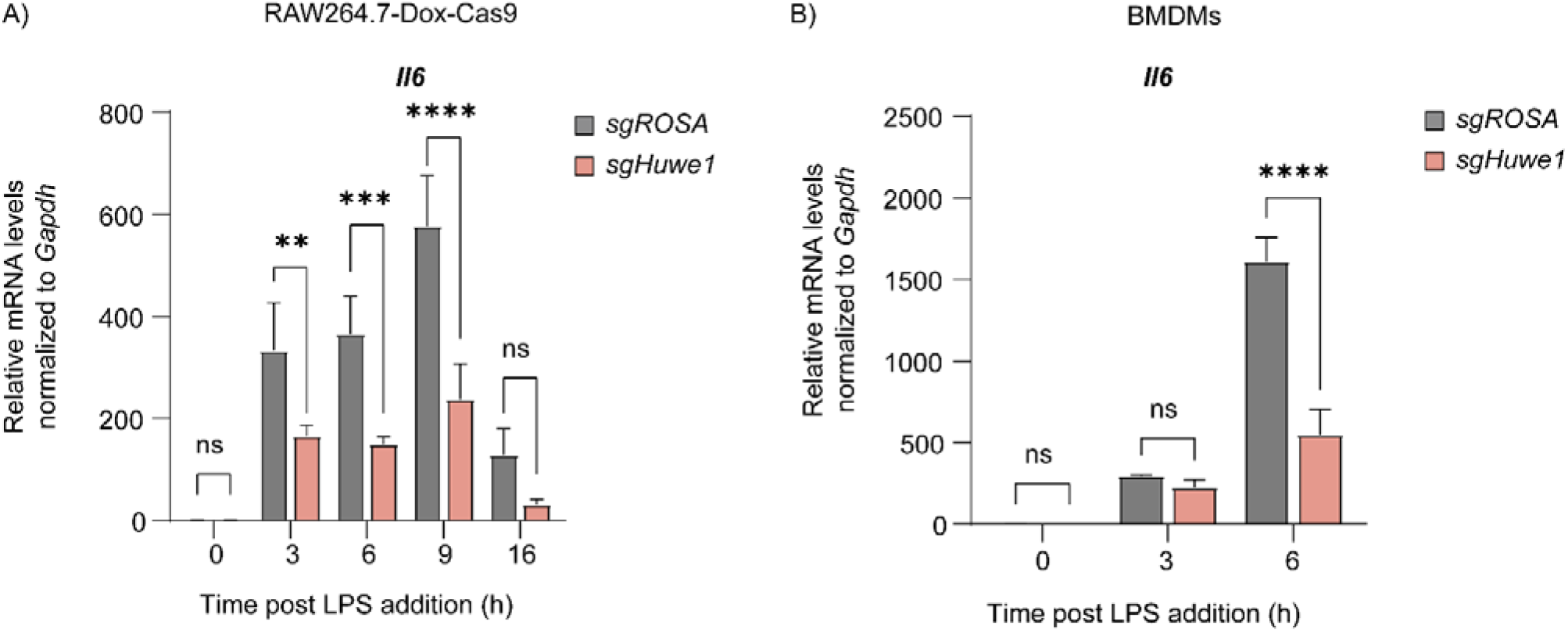
TTP mRNA targets are dysregulated upon *Huwe1* depletion. **A)** RAW264.7-Dox-Cas9 cells expressing sg*ROSA* or sg*Huwe1* were treated with Dox for 5 days to induce Cas9. Cells were incubated with LPS for the indicated time points (h). *Il6* mRNA levels were measured by RT-qPCR and normalized to *Gapdh*. Data represent the mean and s.d.; n = 3. \**p* ≤ 0.05; \*\**p* ≤ 0.01; \*\*\**p* ≤ 0.001; \*\*\*\**p* ≤ 0.0001. Two-way ANOVA was performed. **B)** sg*ROSA*- or sg*Huwe1*-expressing BMDMs were treated with LPS as indicated. *Gapdh*-normalized *Il6* mRNA levels were measured by RT-qPCR. Data represent the mean and s.d.; n = 2. \*\*\*\**p* ≤ 0.0001. Two-way ANOVA was performed.

LPS stimulation transcriptionally induces *Tnf* and *Il6*, and in parallel influences TTP protein stability. Loss of *Huwe1* resulted in significantly decreased concentrations of *Tnf* and *Il6* transcripts at 3 to 16 h. post-stimulation (Fig. EV3A and 3A/B), consistent with increased TTP protein levels in *Huwe1* knock-out cells (Fig. 2C). These differences do not stem from altered macrophage differentiation from bone marrow, as no differences in F4/80 surface expression were measured between sg*ROSA* and sg*Huwe1* BMDMs (Fig. EV3B). Instead, Actinomycin D mRNA chase experiments indicated that the decreased levels of pro-inflammatory mRNAs are consistent with a 57% decrease in *Tnf* mRNA stability (Fig. EV3C). Moreover, targeting of *Zfp36* in *Huwe1*-deficient cells partially rescued *Tnf* mRNA concentrations (Fig. EV3D), indicating that the effects of *Huwe1* loss on *Tnf* mRNA levels are at least in part TTP-dependent.

Taken together, these data show that loss of *Huwe1* increases the bio-active pool of cellular TTP, resulting in enhanced turn-over of TTP target mRNAs encoding pro-inflammatory mediators.

### HUWE1 regulates TTP phosphorylation and its increase is responsible for increased TTP stability

Since HUWE1 is a ubiquitin E3 ligase and was identified as a regulator of TTP protein stability by genetic means, we reasoned that the effects from its ablation on TTP could be direct through complex formation and ubiquitination of TTP, or indirect by influencing the activity or abundance of proteins that regulate TTP. Neither co-IP, nor TurboID proximity labeling assays identified complex formation between TTP and HUWE1 in cells (Appendix Fig. S3 and S4).

This suggested that the effects of HUWE1 on TTP may be indirect, although direct ubiquitination of TTP by HUWE1 cannot be ruled out as their interaction may have been too transient to detect in our assays. Attempts to address direct TTP ubiquitination by HUWE1, or any of the other E3 ligases identified in the genetic screen (Fig. 2B and EV2A-B; VHL, UBE3C, and the Cullin adapters Elongin B/C) were hindered by the inability to purify sufficient amounts of recombinant TTP protein.

Since TTP stability is regulated for an important part through phosphorylation by the stress kinase p38-MK2 axis and to a lesser extent ERK (Brook *et al*, 2006; Ronkina *et al*, 2019; Deleault *et al*, 2008), we set out to determine whether *Huwe1* ablation would alter TTP levels indirectly by affecting the cellular concentrations or activity of these kinases. Data from *Huwe1* knock-out cells indicated that the effect of *Huwe1* loss on TTP stability was predominantly at time points after the initial two hours of LPS stimulation (Fig. 2E) during which TTP dephosphorylation of S52/S178 happens, resulting in its degradation (Sedlyarov *et al*, 2016; Kratochvill *et al*, 2011). Consistent with this finding, *Huwe1* ablation did not significantly alter the total protein levels or change the early phosphorylation/activation kinetics of stress kinases p38, MK2, ERK, and JNK between 0-60 min. post-LPS treatment (Fig. EV4A-D).

In contrast, ablation of *Huwe1* strongly increased endogenous TTP levels upon its induction by LPS at all measured later time points from 2 to 16 h. post-stimulation (Fig. 2E and 4A). In the same lysates, total and activated/phosphorylated levels of p38, MK2, ERK and JNK were determined.

The total levels of all four kinases varied slightly between the different time points post-LPS stimulation, yet these differences were independent of the targeted locus (*ROSA* or *Huwe1*). In contrast, *Huwe1*-targeting consistently increased the activated phosphorylated forms of all kinases at 2 h. post-stimulation (Fig. 4A-D). While activated p38 and ERK levels in *Huwe1*-targeted cells were comparable to the *ROSA*-targeted control, or even lower, at later time points (Fig. 4A-C), MK2 and JNK activation was increased for prolonged times, up to 6 h. post-stimulation (Fig. 4B-D). Importantly, in the absence of LPS stimulation, *Huwe1* knock-out did not affect, or even decreased, baseline phosphorylated levels of all kinases (Fig. 4A-D). Moreover, *Huwe1* ablation in the presence of LPS did not induce unrelated stress responses, such as p53 activation (Fig. EV4E), indicating that loss of *Huwe1* does not induce a general stress response in the cell, but is specific for pro-inflammatory cellular conditions mediated by LPS.

**Figure 4.**
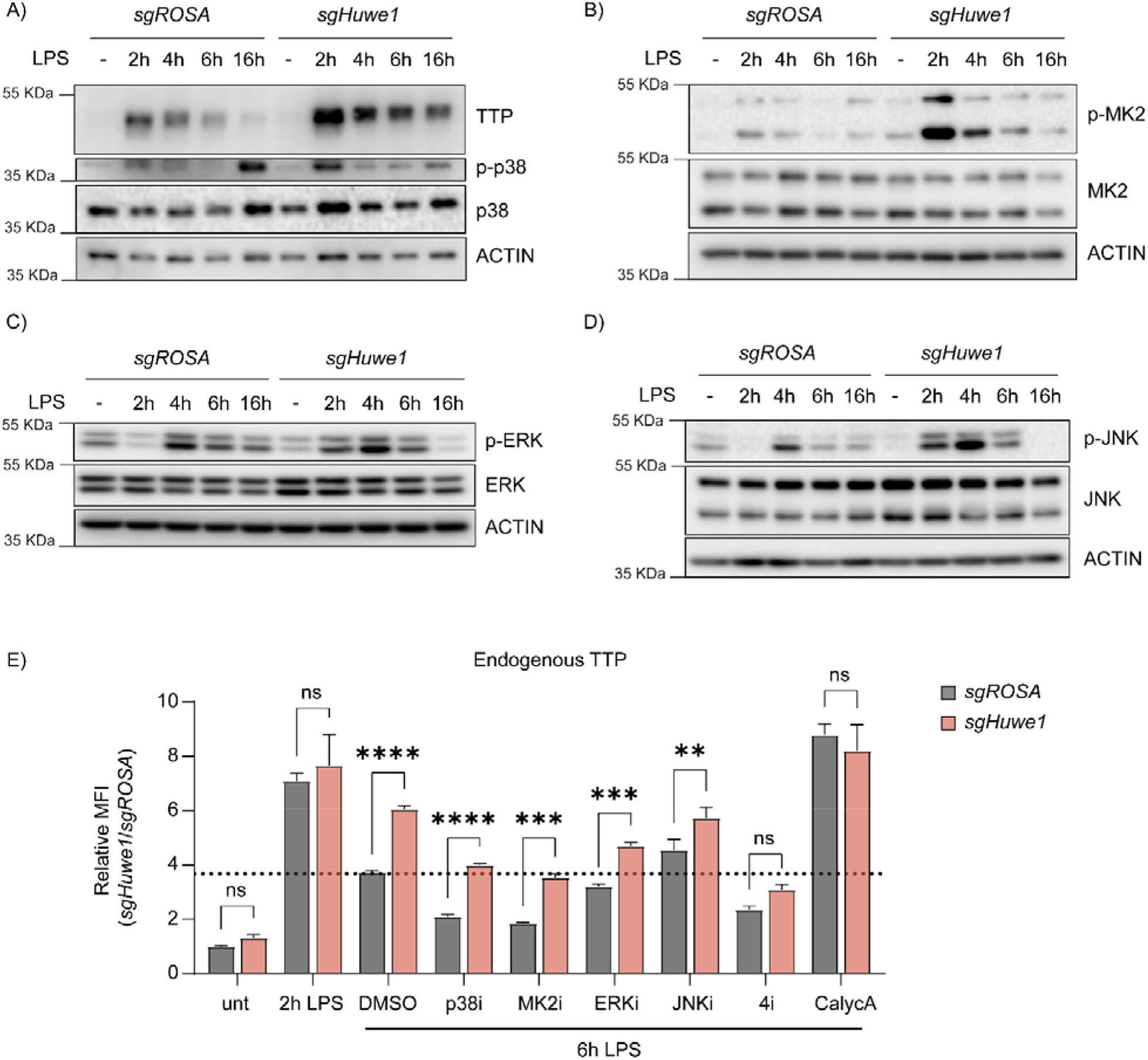
HUWE1 regulates TTP phosphorylation status, and thereby its TTP stability. **A-D)** RAW264.7-Dox-Cas9 cells expressing sg*ROSA* or sg*Huwe1* were treated with Dox for 5 days to induce Cas9. Cells were incubated with LPS for the indicated time points (h). Phosphorylation of **A)** p38, **B)** MK2, **C)** ERK, and **D)** JNK was determined by western blot. **E)** sg*ROSA-* or sg*Huwe1-*RAW264.7-Dox-Cas9 cells were treated with LPS or left untreated. LPS-treated cells were incubated with p38i, MK2i, ERKi, JNKi, or PP1/2 inhibitor Calyculin A (CalycA). TTP levels were analyzed by flow cytometry and normalized MFI plotted. Data represent the mean and s.d.; n = 3. \**p* ≤ 0.05; \*\**p* ≤ 0.01; \*\*\**p* ≤ 0.001; \*\*\*\**p* ≤ 0.0001. Two-way ANOVA was performed. Dotted horizontal line indicates TTP abundance in the DMSO control at 6 h. post-LPS treatment.

Based on the increased levels of phosphorylated TTP (Fig. 2C) and stress kinase activation in *Huwe1* knock-out cells, we hypothesized that phosphorylation of TTP by some or all of the four deregulated kinases could be responsible for the increase in TTP protein stability. In particular p38 and its downstream target MK2 were prime candidates, given their importance in LPS-induced TTP stabilization through phosphorylation of S52 and S172.

To investigate whether the increased activity of the four individual kinases in *Huwe1*-targeted cells was causative for the increased TTP stability, a mixed genetic/inhibitor epistasis experiment was performed. To this end, endogenous TTP levels were assessed by intra-cellular staining in *ROSA-* or *Huwe1-*targeted cells, which were additionally treated with individual inhibitors of p38, MK2, ERK, JNK, or a combination of all four inhibitors (Fig. 4E).

Consistent with our other results, *Huwe1* knock-out increased TTP protein levels in 6 h. LPS-treated DMSO control cells (Fig. 4E; sample set 3). In line with previous reports that the p38-MK2 axis is an important determinant of TTP stability, treatment of sg*ROSA*-targeted cells with p38 or MK2 inhibitors significantly decreased TTP levels (Fig. 4E; sample sets 4 and 5). However, simultaneous *Huwe1* knock-out still elevated TTP levels in the presence of these individual inhibitors, indicating that either HUWE1 does not affect TTP through the p38-MK2 axis, or that there are compensatory mechanisms affecting TTP stability in the absence of p38-MK2 kinase activity.

Consistent with previous findings of a minor effect of ERK activity, and no effect of JNK on TTP stabilization (Deleault *et al*, 2008), ERK or JNK inhibition did not influence TTP levels (Fig. 4E; sample sets 6 and 7). Moreover, the levels of TTP in the presence of either ERK or JNK inhibitors were still increased upon *Huwe1* knock-out, indicating that the activity of neither of these individual kinases alone is required for the HUWE1-dependent effect on TTP.

We hypothesized that the deregulated increase in activity of multiple of these four stress kinases could in a partially functionally compensatory manner contribute to elevated TTP phosphorylation (Fig. 4A-D) and stability. Indeed, inhibition of all four kinases (4i) simultaneously rendered TTP highly unstable as expected, yet in contrast to the single kinase inhibitors, this was no longer affected by *Huwe1* knock-out (Fig. 4E; sample set 8). From these data, we concluded that the HUWE1 effect on TTP stability is dependent on the activity of multiple stress kinases.

Together, these results indicate that HUWE1 is important for curtailing TTP phosphorylation. We reasoned that the increased TTP phosphorylation in *Huwe1* knock-out cells could stem from either increased phosphorylation by the stress kinases, and/or decreased dephosphorylation.

Since MK2/p38, ERK, and JNK are activated/phosphorylated through independent cellular pathways, yet inactivated/dephosphorylated by the same phosphatases as TTP itself (PP1/2) (Nguyen & Shiozaki, 1999; Kruse *et al*, 2020; Takekawa, 1998, 2000; Warmka *et al*, 2001), we reasoned that it was most likely that HUWE1 may be important to regulate PP1/2 activity or its cellular concentrations. Therefore, we hypothesized that decreased PP1/2 output in *Huwe1* knock-out cells could prolong TTP phosphorylation by: **i)** diminishing direct TTP dephosphorylation by PP1/2, and **ii)** indirectly prolonging stress kinases activation as a consequence of their diminished dephosphorylation by PP1/2.

To test this hypothesis, *sgROSA* or *sgHuwe1* cells were treated with LPS for 6 h, and from 2 h. onward, co-incubated with PP1/2 inhibitor Calyculin A (Fig. 4E). As expected, preventing dephosphorylation by this inhibitor stabilized TTP, and prevented TTP degradation by 6 h. of LPS treatment (Fig. 4E; compare sg*ROSA* 6 h. LPS with sg*ROSA* 6 h. LPS + CalycA; sample set 9). In contrast to sg*Huwe1* samples treated for 6 h. with LPS (in which TTP protein levels were increased), treatment with Calyculin A rescued the *Huwe1* knock-out-dependent increase in TTP protein concentrations (Fig. 4E; compare sg*Huwe1* 6 h. LPS with sg*Huwe1* 6 h. LPS + CalycA).

From these results, we conclude that in the absence of HUWE1, decreased cellular output of PP1/2 may prolong stress kinase activation. Increased kinase activity and decreased dephosposphorylation of TTP by PP1/2 consequently increases TTP phosphorylation, thereby stabilizing it.

### HUWE1 controls only a small fraction of proteasome targets, and regulates the stability of TTP paralog ZFP36L1

HUWE1 has been shown to associate with proteasomes (Besche *et al*, 2009), the biological significance of which has remained elusive. We reasoned that HUWE1 might be important for proteasome activity, and its ablation cause a general impaired degradation of proteasome targets such as TTP. To investigate whether this was the case, we compared the proteomes of LPS-stimulated RAW264.7-Dox-Cas9 cells in which we targeted either *Huwe1* or proteasome core particle component *Psmb7* by label-free mass-spectrometry.

As expected, *Psmb7* targeting altered the abundance of a large number of proteins, many of which are known targets of proteasomal degradation (Fig. EV5A and Table EV2). In contrast, *Huwe1* ablation significantly changed the concentrations of only a select number of proteins (Fig. 5A and S5B). In line with expectations of an E3 ligase, HUWE1 targets showed a trend of also being increased in *Psmb7* knock-out cells, and *vice versa* (Fig. 5A and EV5A/B). However, there was no clear correlation between the most affected proteins in the two genotypes, indicating that HUWE1 is likely not essential for proteasome function in cells, and that the increase of TTP in *Huwe1* knock-out cells is unlikely to have resulted from diminished overall proteasome activity. Among the differentially regulated proteins were factors previously identified as HUWE1 targets (Cassidy *et al*, 2020; Xu *et al*, 2016; Thompson *et al*, 2014), including GRB2, CHEK1, and CDC34 (Fig. EV5C).

**Figure 5.**
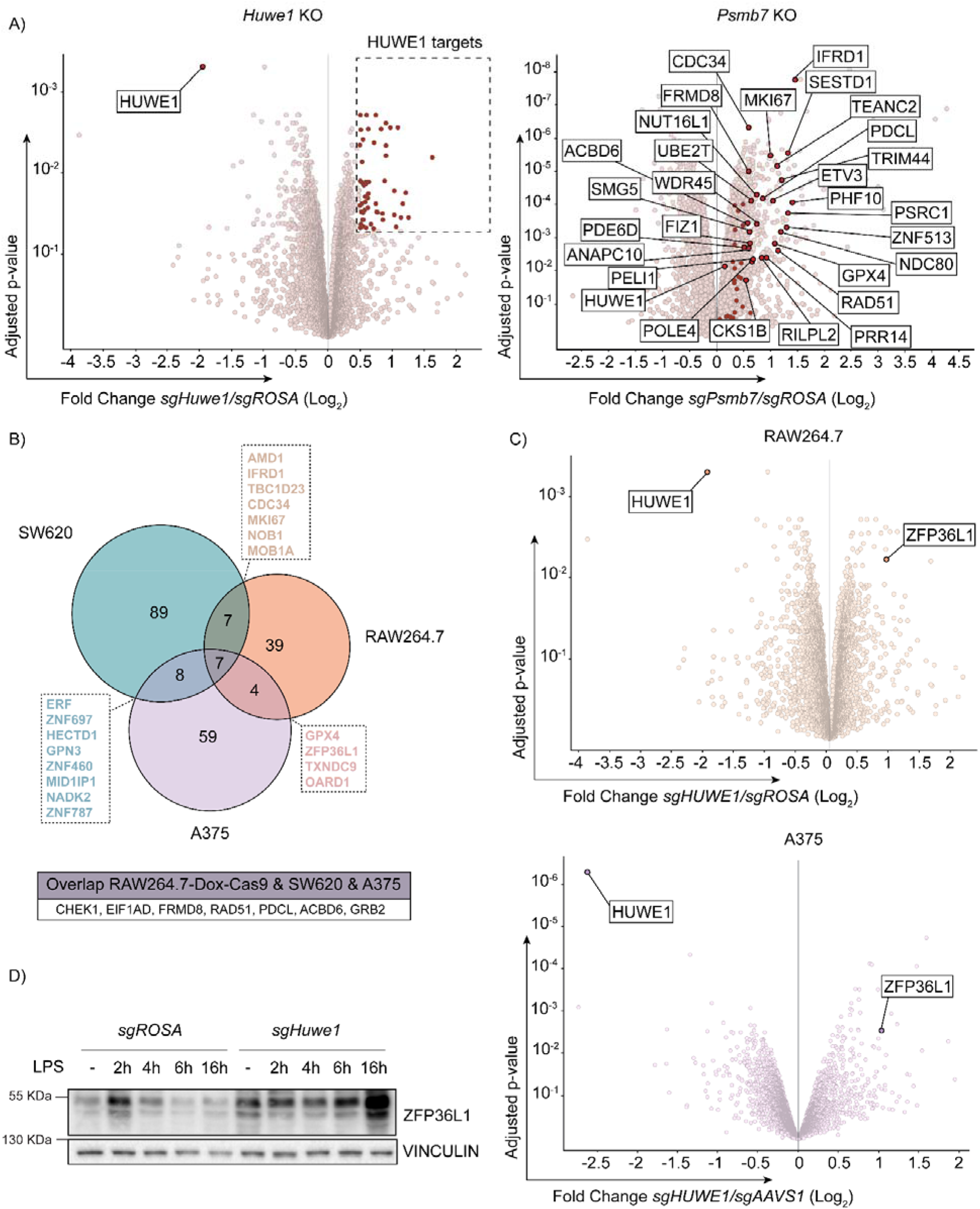
TTP family member ZFP36L1 is stabilized upon HUWE1 knockout in human and mouse. **A)** RAW264.7-Dox-Cas9 expressing sg*ROSA*, sg*Huwe1* or sg*Psmb7* were treated with Dox for 3 days to induce Cas9. Proteome changes were assessed by quantitative mass spectrometry. Proteins classified as HUWE1 targets are highlighted in red. Shared HUWE1 and proteasome targets are labelled in the *Psmb7* knock-out volcano plot. (adjusted p-value ≤ 0.05 and Fold Change (Log_2_) ≥ 0.5; n = 3). **B)** Venn diagram showing the overlap between proteome changes of *Huwe1*-targeted RAW264.7, A375, and SW620 cell lines. Shared targets are listed (adjusted p-value ≤ 0.05 and Fold Change (Log_2_) ≥ 0.5; n = 3). **C)** Volcano plots representing proteome changes of *Huwe1*- and *AAVS1*/*ROSA*-targeted A375 human melanoma cells and RAW264.7-Dox-Cas9 cells (adjusted p-value ≤ 0.05 and Fold Change (Log_2_) ≥ 0.5; n = 3). The shared HUWE1 target ZFP36L1 is highlighted. **D)** sg*ROSA* or sg*Huwe1* knockout RAW264.7-Dox-Cas9 cells were treated with Dox for 5 days to induce Cas9, followed by LPS treatment for the indicated times (h). ZFP36L1 protein levels were determined by western blot.

Our data so far indicated that HUWE1 is important for proper regulation of TTP phosphorylation, and that in its absence the equilibrium shifted to a hyper-phosphorylated state (Fig. 4), dependent on a decrease in phosphatase activity, and an increase in stress kinase activity, without major effects on their protein levels.

To further assess in an unbiased manner whether *Huwe1* deficiency would affect MAPK or PP1/2 protein levels, we extended our proteome mass-spectrometry for two additional human cell lines (A375 and SW620). Consistent with our previous findings (Fig. 4), *Huwe1* ablation did not substantially or consistently affect the protein levels of detected MAPK or PP1/2 subunits in the three cell lines (Fig. EV5D/E). In line with the data presented above (Fig. 4), this suggests that the hyper-phosphorylated TTP state in *Huwe1*-targeted cells does not result from changes in MAPK or PP1/A protein levels.

Previous studies have indicated that HUWE1 can target broader classes of cellular substrates (Hunkeler *et al*, 2021; Grabarczyk *et al*, 2021), but that the targeted proteins may be cell type specific to some degree. Analysis of proteome changes in the three different cell lines identified seven proteins that were consistently increased in all *Huwe1*-targeted cell lines (Fig. 5B). Moreover, the protein concentration of other proteins was changed in only two of the three cell lines, whereas it was not detected in the third (Fig. 5B).

We reasoned that any of these common deregulated proteins in *Huwe1* knock-out cells could contribute to the TTP hyper-phosphorylation/stabilization phenotype. However, analysis of overlap between factors that regulate TTP abundance identified in the genetic screen (Fig. 2B and EV2A), and proteins deregulated by *Huwe1*-ablation did not identify any overlap, suggesting that these *Huwe1*-regulated proteins are unlikely to drive the effect on TTP protein stability.

Importantly, proteome measurements by mass-spectrometry are limited to detection of only reasonably abundant proteins. Even after LPS stimulation, no TTP peptides were identified in any of the three analyzed cell lines (Fig. 5B), indicating that its absolute intra-cellular concentrations in these cells are too low to be detected by this method. In contrast, peptides of its paralog ZNF36L1 were readily identified in RAW264.7 and A375 cells and among the most increased proteins identified in *Huwe1*-targeted cells (Fig. 5C). In line with this observation, independent western blot analysis of ZFP36L1 in cell lysates from *Huwe1*-deficient RAW264.7 cells cells showed that ZFP36L1 protein levels were increased in *Huwe1* knock-out cells (Fig. 5D).

Collectively, these findings indicate that *Huwe1* ablation does not alter MAPK or PP1/2 protein levels, but that rather their differential activation alters TTP phosphorylation and stability. Moreover, ZFP36L1 abundance is regulated by HUWE1, akin to its closest related family member -TTP-, indicating that they could be regulated by HUWE1 in a conserved manner.

### Residues in the TTP 234-278 region are important for its stability

Lysines are exclusively located in the TTP zinc finger domain, and our data indicate that this is the site of poly-ubiquitination (Fig. EV1B). In line with this notion, upon mutation of the five lysine residues in its zinc finger domain (KtoR), TTP accumulated as a stable, phosphorylated species (Fig. 6A-B and EV6A-B). Moreover, this mutant was no longer affected by *HUWE1* loss, indicating that the effects of HUWE1 on TTP stability are dependent on ubiquitination in the zinc finger domain.

**Figure 6.**
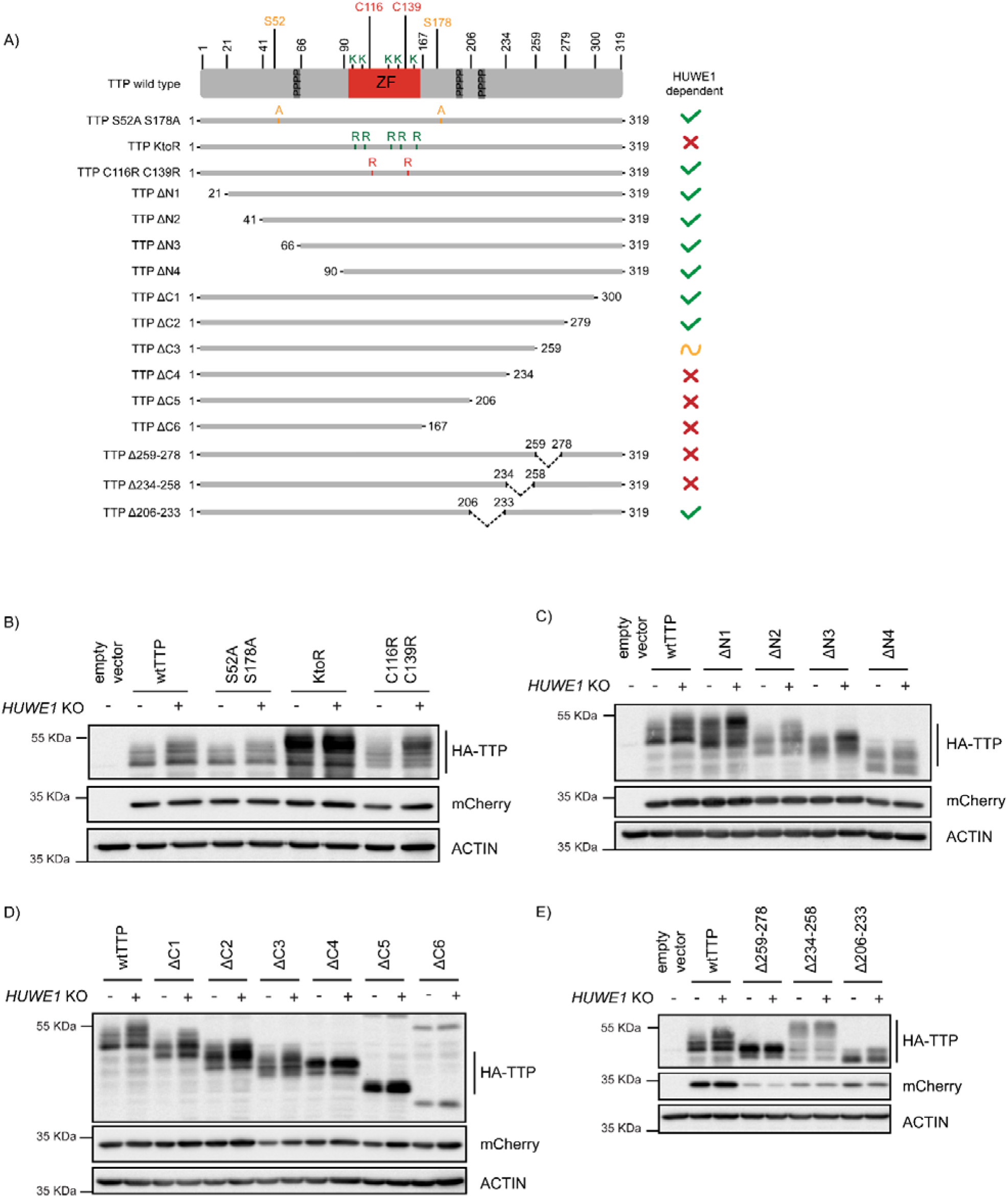
**A)** Schematic representation of 3xHA-TTP mutants. Colors denote amino acid substitutions. ZF indicates the zinc finger domain, and the three tetraprolin motifs are presented as dark grey boxes. **B)-E)** sg*AAVS1- and* sg*HUWE1*-depleted HEK293T cells were transfected with the indicated mutants, and 3xHA-TTP stability was determined by western blot. mCherry is expressed as a stable internal control through a P2A site.

We reasoned that the E3 ligase ubiquitinating TTP at that site could bind TTP in its folded zinc finger domain, or that the mRNA-engaged TTP pool could be the predominant HUWE1-dependent target. Therefore, we addressed whether a TTP mutant with a disrupted zinc finger domain (C116R, C139R) would be stabilized. Zinc finger domain disruption did not accumulate at higher steady-state levels than its wtTTP counter-part, and was still increased upon *HUWE1* loss (Fig. 6B and EV6B), demonstrating that neither recognition of the folded zinc finger domain structure by an E3 ligase, nor TTP functionality are required for the HUWE1 effects.

Our data support a role of HUWE1 in determining TTP phosphorylation, and thereby its stability. Therefore, we next analyzed whether phosphorylation of the two best-characterized TTP residues in this context (S52, S178) are important for HUWE1 effects. As for the zinc finger domain mutant, an (S52A/S178A) TTP mutant was still stabilized by *HUWE1* loss (Fig. 6B and EV6B). This indicated that while phosphorylation of these residues importantly controls TTP stability, HUWE1 effects are likely independent of these phospho-residues.

Together, these data from full-length TTP point mutants suggest that an E3 ligase likely binds TTP outside of its zinc finger domain, but ubiquitinates it on lysines inside the zinc finger domain. Moreover, we concluded that HUWE1 regulation of TTP stability and phosphorylation is independent of the MK2-stabilized S52/S178 residues. This is consistent with the finding that TTP levels were still increased in *Huwe1* knock-out cells treated with inhibitors of the p38 and MK2 kinases phosphorylating these two sites (Fig. 4E; subsets 4-5). Lastly, these data indicate that TTP does not require an intact zinc finger domain for stability, suggesting that TTP engagement with target mRNAs is likely not a prerequisite for HUWE1-dependent stability regulation.

Next, we set out to determine which part of TTP regulates its HUWE1-dependent phosphorylation and stability. To this end, progressive N- and C-terminal TTP deletion mutants (Fig. 6A) were analyzed in cells for their steady-state concentrations, phosphorylation, and sensitivity to *HUWE1* ablation. Since TTP is predicted to be mostly disordered outside of its zinc finger domain (https://alphafold.ebi.ac.uk/entry/P22893), we reasoned that the effects of the truncations on overall protein structure would be limited.

N-terminal deletions did not affect TTP protein levels, its phosphorylation, or the effect of *HUWE1* loss (Fig. 6C and EV6C; deletions ΔN1-4), indicating that HUWE1 does not influence TTP stability through residues N-terminal of the zinc finger domain. Likewise, the two most C-terminal deletions did also not affect TTP stability (Fig. 6D and EV6D; ΔC1-2). In contrast, further truncation of the C-terminus rendered mutant ΔC3 (259-278 region) less sensitive to *HUWE1* loss, yet retained its hetrogeneous size distribution for phosphorylated species. Further deletion of the 234-258 region in the ΔC4 mutant strongly stabilized TTP at a homogeneously phosphorylated size, and rendered it insensitive to *HUWE1* knock-out (Fig. 6D and EV6D). Likewise, the ΔC5 and ΔC6 mutants were insensitive to *HUWE1* knock-out, but accumulated as unphosphorylated TTP species. Together, these data indicate that the 234-278 region (Fig. 6D and EV6D) is important for HUWE1-dependent regulation of TTP stability, and its phosphorylation status.

Since the TTP-ΔC3 mutant was stabilized in a HUWE1-insensitive manner (Fig. 6D and EV6D), we reasoned that this region (259-278) could be important for proteasomal targeting (e.g. an E3 ligase binding site). In contrast, the TTP-ΔC4 mutant accumulated as a lower MW homogenously phosphorylated TTP species (Fig. 6D and EV6D), suggesting that the 234-258 region regulates TTP stability by affecting its phosphorylation status (e.g. a phosphatase binding site).

To test these possibilities, we analyzed mutants in which either only the 259-278 or 234-258 regions were deleted, while retaining the rest of the protein (Fig. 6A). Consistent with the data from Fig. 6B, a TTP mutant only lacking the 259-278 region was strongly stabilized, accumulated predominantly as a relatively homogeneous phosphorylated species, and was insensitive to *HUWE1 knock-out* (Fig. 6E and EV6E). Moreover, deletion of the 234-258 resulted in TTP hyper-phosphorylation (Fig. 6E and EV6E), consistent with the idea of it being important for phosphatase binding.

Together, these data indicate that the TTP ΔC3-specific region (259-278) is consistent with a possible binding site for an E3 ligase, whereas the ΔC4-specific region (234-258) is a likely interaction site of a phosphatase (Fig. 7A). Importantly, the ΔC5- and ΔC6-mutants were stabilized, yet not hyperphosphorylated (Fig. 6B). This suggests that the phosphorylated residues contributing to TTP stabilization in *HUWE1* knock-out cells are likely in the possible E3 ligase binding site in the TTP ΔC3-specific region (259-278) (Fig. 7A).

**Figure 7.**
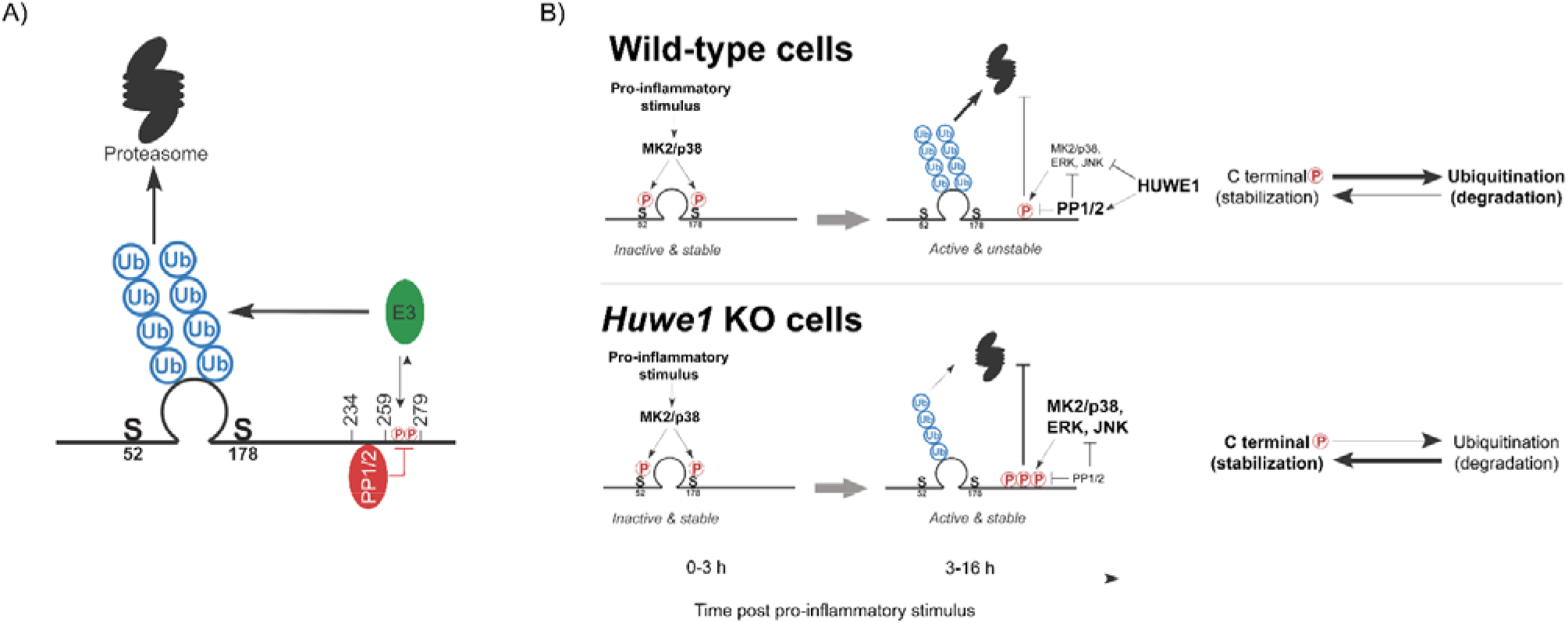
**A)** Model indicating the TTP regions in its C-terminus speculated to recruit PP1/2 and an E3 ligase, which ubiquitinates the zinc finger domain. **B)** Model of TTP stability regulation through phosphorylation in wild-type cells and *Huwe1*-deficient cells.

In summary, we provide evidence for ubiquitin-dependent proteasomal degradation as a key regulatory mechanism for TTP protein abundance in cells. A genetic screen identified HUWE1 as a strong regulator of TTP proteasomal turn-over. In the absence of *Huwe1*, TTP is heavily phosphorylated and stabilized, which is dependent on ubiquitination sites in the TTP zinc finger domain, and phosphorylation in the 259-278 region. We propose that this region in its unphosphorylated form is also a likely binding site for an E3 ligase directing TTP ubiquitination and degradation (Fig. 7A). Moreover, the adjacent 234-258 region is consistent with an interaction site for the main TTP phosphatases (PP1/2) (Fig. 7A).

We propose a model in which HUWE1 under physiological conditions curtails stress kinase activation, thereby limiting their stabilizing effects on TTP (Fig. 7B). However, in the absence of *Huwe1*, the collective activity increase of these stress kinases results in TTP hyper-phosphorylation in the 259-278 region, increased TTP stability, and decreased pro-inflammatory output. Since we found that TTP phosphorylation is inversely correlated with ubiquitination and degradation, we speculate that phosphorylation in this region could prevent E3 ligase binding (Fig. 7A).

## Discussion

Previous studies have predicted that TTP is disordered outside of its zinc finger domain, and showed that these unstructured regions contribute to its rapid proteasomal turn-over (Ross *et al*, 2015; Ngoc *et al*, 2014). Protein disorder is often associated with proteasomal turn-over, as these regions often contain degrons, accessible ubiquitination sites, or provide an initiation side for threading into the proteasome and initiating unfolding and translocation into the catalytic chamber (Aufderheide *et al*, 2015; van der Lee *et al*, 2014). However, previous work did not identify TTP poly-ubiquitination, and showed that incubation of TTP with purified 20S proteasomes -which lack the Ub-receptor containing 19S regulatory particle-, were sufficient to degrade TTP (Ngoc *et al*, 2014).

It has been reported that oxidized, unfolded proteins could be directly degraded by 20S proteasomes (Davies, 2001; Inai & Nishikimi, 2002). However, the prevailing notion is that association of a regulatory particle is critical to open access to the catalytic chamber (Coux *et al*, 1996; Davies, 2001; Driscoll & Goldberg, 1990; Eytan *et al*, 1989), and efficient substrate degradation in cells. Here, we demonstrate robust poly-ubiquitination of TTP in denaturing RAW264.7 lysates, indicating that these poly-ubiquitin chains are covalently attached to TTP, and do not interact through a putative ubiquitin-interaction domain. Moreover, we show that mutation of lysines in the zinc finger domain stabilized TTP, and that an inhibitor preventing *de novo* ubiquitination in cells, stabilized TTP. These data demonstrate that poly-ubiquitination of TTP is essential for its degradation, which is likely mediated by 26S proteasomes.

Most proteins with disordered regions will eventually be degraded in *in vitro* reactions containing high concentrations of 20S proteasomes (Lu *et al*, 2015; Liu *et al*, 2003). This could explain the previous finding of ubiquitin-independent TTP degradation *in vitro* (Ngoc *et al*, 2014). Future comparisons of TTP degradation kinetics of ubiquitinated and non-ubiquitinated forms in the presence of 26S proteasomes will be important to further address this issue.

Several E3 ligases to putatively poly-ubiquitinate TTP were identified in our genetic screen (Fig. 2): HUWE1, VHL, UBE3C, and the Cullin adapters Elongin B/C. *Huwe1* ablation most robustly stabilized TTP, based on which we hypothesized that HUWE1 may directly poly-ubiquitinate TTP. However, multiple independent techniques, including TurboID proximity labeling and co-IPs, failed to identify an interaction between TTP and HUWE1, which suggested that it may instead indirectly influence TTP stability.

At this point, we cannot rule out that HUWE1 directly poly-ubiquitinates TTP, resulting in its proteasomal degradation. Alternatively, one or more of the other identified E3 ligases could contribute to direct TTP ubiquitination. If indeed the other E3 ligases contribute to TTP ubiquitination, the fact that their knock-out phenotype is substantially less than that of *Huwe1* loss (Fig. EV2B) may suggest that multiple of them could be functionally redundant. Irrespective of potential direct TTP ubiquitination by HUWE1, multiple lines of evidence point towards a strong contribution of HUWE1-dependent differential TTP phosphorylation as an indirect means to control TTP stability and functional activity. This is consistent with our finding that TTP phosphorylation and ubiquitination appear to be inversely correlated, as the TTP KtoR mutant accumulates as a hyper-phosphorylated species (Fig. 6B).

Published data support a model in which TTP upon translation is initially phosphorylated by the p38/MK2 kinase axis on residues S52 and S178 (mTTP numbering), resulting in its stabilization, yet repressing TTP function (Deleault *et al*, 2008; Hitti *et al*, 2006; Ross *et al*, 2015; Kratochvill *et al*, 2011). At later stages, diminishing p38/MK2 kinase activity is thought to shift the equilibrium to dephosphorylation of TTP at these residues, rendering it active, but unstable. Under these conditions TTP is rapidly turned over by the proteasome (Ross *et al*, 2015; Deleault *et al*, 2008).

Consistent with these data, we also found that an S52A/S178A mutant is unstable in the absence and presence of LPS stimulation (Fig. 6B). Importantly, this mutant was still stabilized in the absence of *Huwe1*, indicating that the stabilizing effects of phosphorylation on S52/S178 are independent of HUWE1. Moreover, it indicates that HUWE1-dependent effects on TTP stability target other sites in TTP. Thus, S52/S178 phosphorylation seems predominantly relevant for TTP stabilization at the early (2-3 h) time points post-LPS-stimulation (Ross *et al*, 2015), whereas HUWE1-dependent effects occur later between 3-16 h) post-LPS stimulation.

In contrast to S52/178, TTP phosphorylation in its C-terminal 259-278 region and its associated stabilization in the absence of *Huwe1*, is paralleled by decreased concentrations of known TTP mRNA targets, which suggests that phosphorylation in this region may not inhibit TTP functional output.

To uncouple LPS-induced *Zfp36* transcription from PTMs influencing TTP protein stability, we complemented experiments using endogenous TTP with exogenously expressed counterparts. In the absence of LPS stimulation, *Huwe1* loss did increase exogenous TTP, albeit rather mildly (Fig. 2C and S2B). In contrast, upon LPS stimulation and stress kinase activation, the effect of *Huwe1* knock-out was much stronger, resulting in strong TTP protein accumulation, which included a substantial fraction of phosphorylated forms (Fig. 2C, 2F, and EV2B). These results suggest that there may be low baseline levels of activated stress kinases in the cell that affect TTP stability in the absence of LPS. However, in contrast to these mild phenotypes, the predominant effects of *Huwe1* loss on TTP protein stability and phosphorylation occur after LPS stimulation, in line with the notion that HUWE1 regulates TTP stability through influencing stress kinase-dependent TTP phosphorylation (Fig. 5 and 6).

*Huwe1* loss increased the activity of multiple stress kinases (Fig. 4A-D) without affecting their total protein levels. The previous findings that **i)** the phosphorylation and activation of these kinases is controlled by PP2A (Nguyen & Shiozaki, 1999; Kruse *et al*, 2020; Takekawa, 1998), and **ii)** the observation that combined inhibition of the stress kinases, or inhibition of PP1/2 activity with Calyculin A, rescued *Huwe1* knock-out effects on TTP (Fig. 4E), suggest that HUWE1 may be important for PP1/2 activity. In line with this notion, decreased PP1/2 activity in *Huwe1* knock-out cells could affect TTP phosphorylation and stability by directly affecting its dephosphorylation, and in parallel maintain high stress kinase activity, which increases TTP phosphorylation even more.

This functional interaction between HUWE1 and the activity of PP1/2 and stress kinases has not been described previously, although it should be noted that HUWE1 has been reported to control the abundance and activity of other kinases and phosphatases (Cassidy *et al*, 2020; Su *et al*, 2021; Jang *et al*, 2014). Our findings broaden the understanding of how HUWE1 may indirectly influence numerous cellular proteins beyond direct recognition as ubiquitination substrates.

Taken together, our data support a model in which HUWE1 is important to maintain PP1/2 cellular output, and curtail stress kinase activation. This in turn limits phosphorylation in the TTP region spanning residues 259-278, allowing for recruitment of HUWE1 itself or another -yet unidentified-E3 ligase to that same region, subsequent poly-ubiquitination on lysine residues in the zinc finger domain, and ultimately proteasomal degradation. Phosphorylation in this region could prevent E3 ligase binding. Although the HUWE1-dependent phosphorylation effect appears to be dependent on the putative E3 ligase binding site (259-278), phosphatase recruitment to the 234-258 region in TTP seemingly controls dephosphorylation of most or all phospho-sites on TTP (Fig. 6C). Since ZFP36L1 abundance is also strongly regulated by HUWE1, this suggests that its C-terminal region is orthologous to TTP 234-278 and how it controls HUWE1-dependent degradation could be conserved across these family members.

## Acknowledgements

We are grateful to Johannes Bock for establishing reagents and methodology that enabled this work. We thank Kitti Csalyi, Johanna Stranner, and Thomas Sauer at the Max Perutz Labs BioOptics FACS Facility for expert support, Markus Hartl and the Max Perutz Labs Mass Spectrometry Facility for mass spectrometry analysis, the IMP/IMBA Protein Biochemistry Core Facility for performing quantitative proteomics, and the Vienna Biocenter Core Facilities (VBCF) for Next Generation Sequencing analysis. We thank Laura Boccuni for expert advice, and the ‘Signaling Mechanisms in Cellular Homeostasis’ doctoral program community, Manuela Baccarini, Thomas Decker, Pavel Kovarik and their labs for their technical expertise and help.

## Author contributions

S.S., A.Sö., A.Sh., I.S., K.H., R.G., R.W.K. performed experiments and analyzed data. M.S. and V.B. established critical reagents and methodology. K.F. performed bioinformatic analyses. A.B., M.S., T.C., P.K, J.Z. provided critical input on experimental designs and data analyses. G.A.V. designed and supervised the study and wrote the manuscript, together with S.S., with feedback from all co-authors.

## Expanded View Figure Legends

**Figure EV1.**
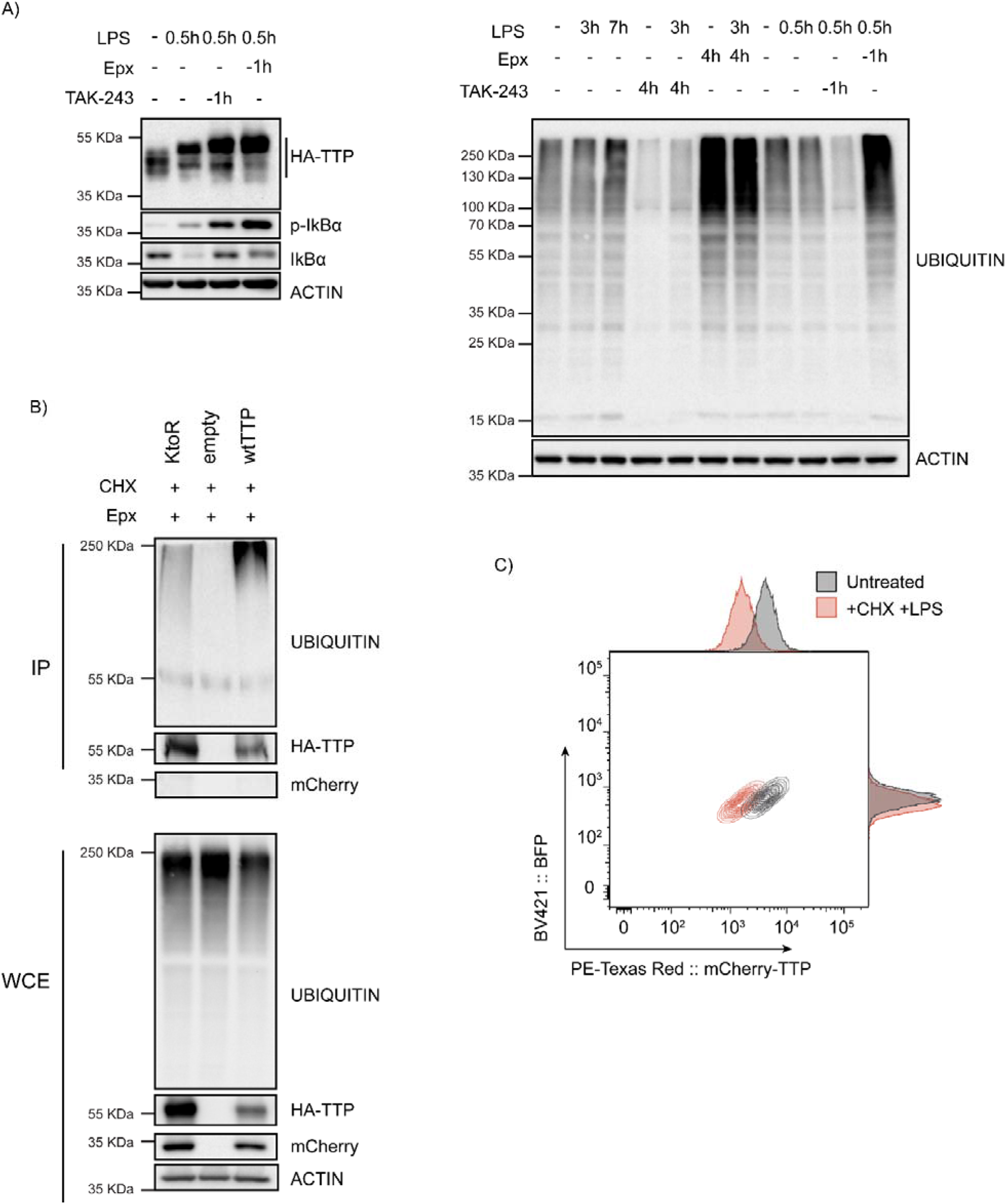
TTP is degraded by the proteasome in a ubiquitin-dependent manner. **A)** 3xHA-TTP-expressing RAW264.7 cells were incubated with LPS or left unstimulated. Cells were then treated with E1 enzyme inhibitor (TAK-234) or the proteasome inhibitor Epoxomicin (Epx). Ubiquitin levels were assessed by western blot. **B)** HEK-293T cells were transiently transfected with plasmids encoding 3xHA-tagged wild-type TTP, a TTP KtoR mutant, or an empty vector. Cells were incubated with the translation inhibitor cycloheximide (CHX) and Epx for 5 h. Immunoprecipitation was carried out using HA antibody, and TTP ubiquination was analysed by western blot. **C)** RAW264.7-Dox-Cas9-mCherry-TTP cells were either left unstimulated or treated with LPS for 5 h. Subsequently, cells were incubated with CHX for 10 h. and analyzed for mCherry-TTP protein levels by flow cytometry.

**Figure EV2.**
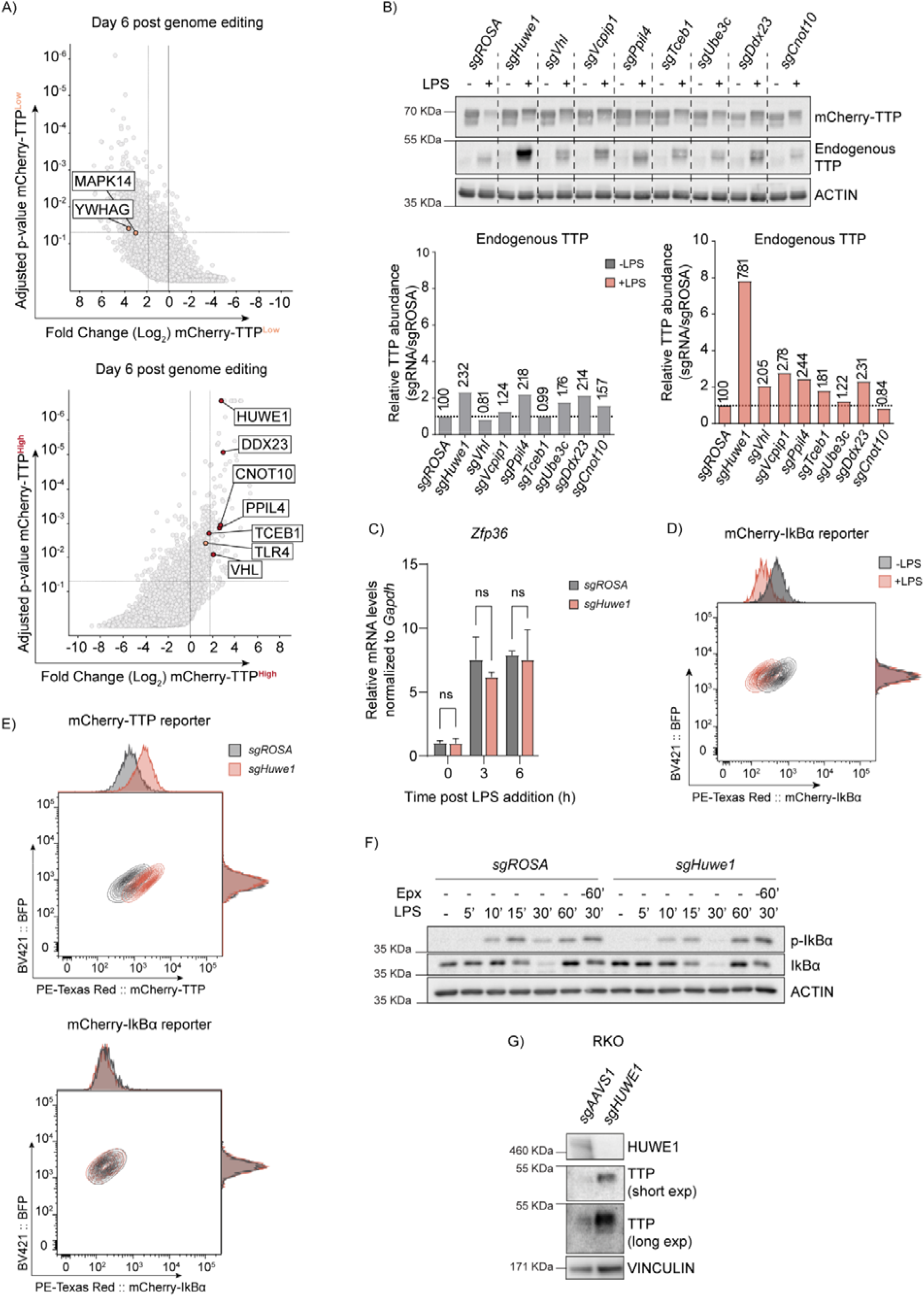
Genome-wide CRISPR-Cas9 knockout screen identified the E3 ligase HUWE1 as a regulator of TTP stability. **A)** Read counts per million in the mCherry-TTP^high^ cells at 6 days after Cas9 induction were compared to those in unsorted cells from the same day, sgRNA enrichment calculated by MAGeCK analysis, and log_2_-fold change and adjusted p-value plotted. Genes enriched in the sorted populations that met the following criteria are indicated in red: a log_2_ fold-change of < 1.8 (mCherry-TTP^low^) or > 1.8 (mCherry-TTP^high^), adjusted p-value < 0.05, not enriched in the matching eBFP2^low^ or eBFP2^high^ sorted cells. **B)** RAW264.7-Dox-Cas9 cells were transduced with lentiviral vectors encoding the indicated sgRNAs. Cas9 was induced for 5 days with Dox. Subsequently, cells were treated with LPS for 16 h, and TTP protein levels were analyzed by western blot. **C)** RAW264.7-Dox-Cas9 cells expressing sg*ROSA* or sg*Huwe1* were treated with Dox for 5 days to induce Cas9. Cells were incubated with LPS for the indicated time points (h). *Zfp36* mRNA levels were measured by RT-qPCR and normalized to *Gapdh*. Data represent the mean and s.d.; n = 3. Two-way ANOVA was performed. **D)** RAW264.7-Dox-Cas9 cells expressing mCherry-IκBα were either left unstimulated or treated with LPS for 2 h, and mCherry-IκBα protein levels analyzed by flow cytometry. **E)** RAW264.7-Dox-Cas9 cells stably expressing mCherry-TTP or mCherry-IκBα fusion proteins were transduced with lentiviral expression constructs encoding sg*ROSA* or sg*Huwe1*. Knock-out was induced for 3 days by the addition of Dox. Cells were stimulated with LPS for 2 h. (mCherry-IκBα) or 16 h. (mCherry-TTP), after which mCherry and eBFP levels were determined by flow cytometry. **F)** RAW264.7-Dox-Cas9 expressing sg*ROSA* or sg*Huwe1* were treated with Dox for 5 days to induce Cas9. Cells were pre-treated with Epx for 60 min. or left unstimulated. Subsequently, cells were incubated with LPS for the indicated times and endogenous IκBα phosphorylation and degradation was analyzed by western blot. **G)** RKO-Dox-Cas9 cells expressing sg*AAVS1* or sg*HUWE1* were stimulated with Dox for 6 days, after which TTP protein levels were determined by western blot.

**Figure EV3.**
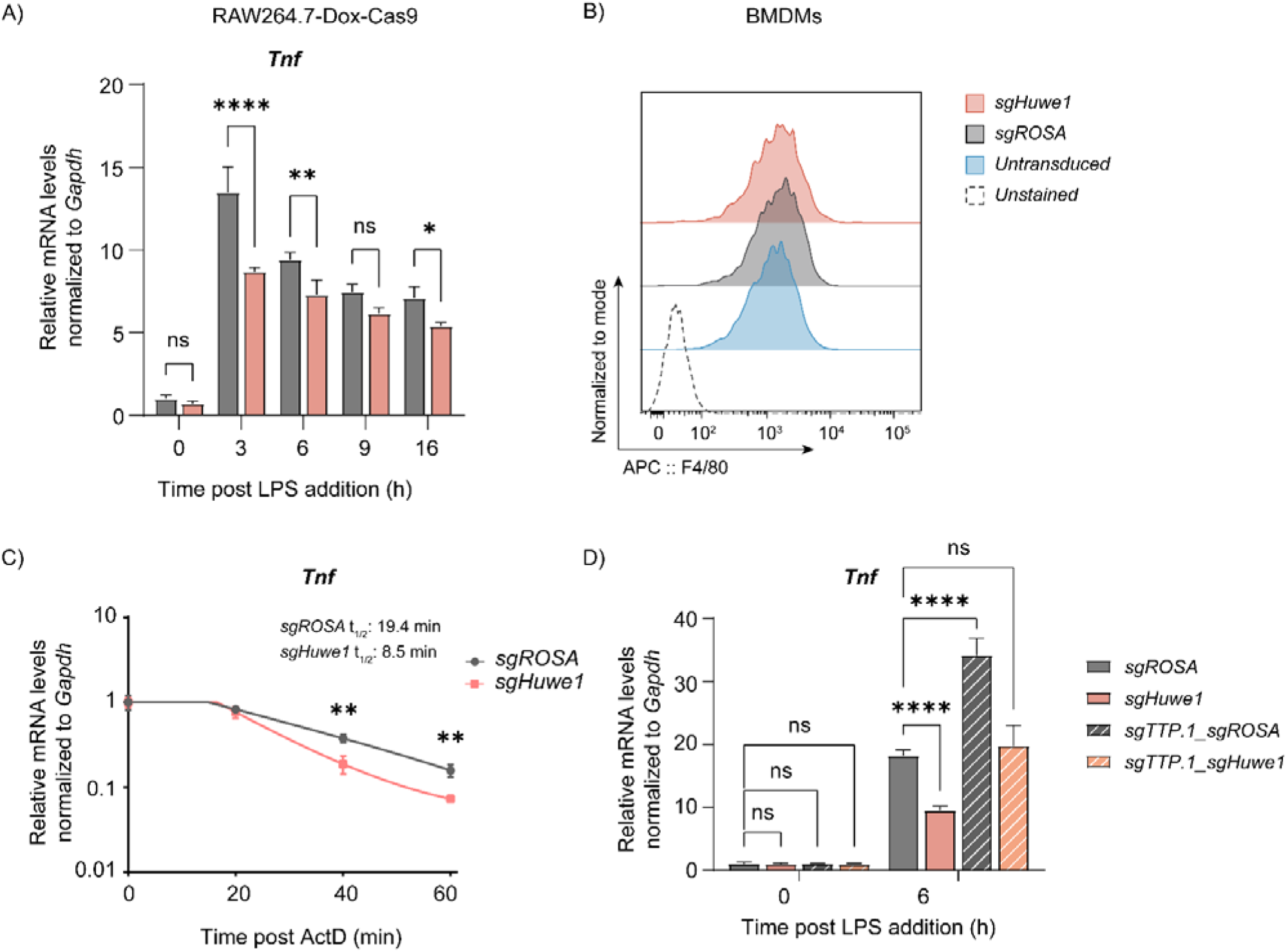
TTP mRNA targets are dysregulated upon *Huwe1* depletion. **A)** RAW264.7-Dox-Cas9 cells expressing sg*ROSA* or sg*Huwe1* were treated with Dox for 5 days to induce Cas9. Cells were incubated with LPS for the indicated time points (h). *Tnf* mRNA levels were measured by RT-qPCR and normalized to *Gapdh*. Data represent the mean and s.d.; n = 3. \**p* ≤ 0.05; \*\**p* ≤ 0.01; \*\*\**p* ≤ 0.001; \*\*\*\**p* ≤ 0.0001. Two-way ANOVA was performed. **B)** Differentiation of sg*ROSA*- or sg*Huwe1*-expressing BMDMs after 7 days of M-CSF treatment was analyzed by F4/80 staining and analysis by flow cytometry. **C)** RAW264.7-Dox-Cas9 cells expressing sg*ROSA* or sg*Huwe1* were treated with Dox for 5 days to induce Cas9. Cells were incubated with LPS for 3 h, after which Actinomycin D (ActD) was added for the indicated times (min)., and *Tnf* mRNA levels were determined by RT-qPCR. Data represent the mean and s.d.; n = 3. \**p* ≤ 0.05; \*\**p* ≤ 0.01; \*\*\**p* ≤ 0.001; \*\*\*\**p* ≤ 0.0001. Unpaired t-test was performed for the 40 min. and 60 min. time point. **D)** RAW264.7-Dox-Cas9 cells expressing sg*ROSA* or sg*Huwe1* single sgRNA vectors, or sg*ROSA-sgTTP* or sg*Huwe1-sgTTP* double sgRNA vectors were incubated with LPS for the indicated times. *Tnf* mRNA expression was determined by RT-qPCR. Data represent the mean and s.d.; n = 3. *p ≤ 0.05; **p ≤ 0.01; ***p ≤ 0.001; ****p ≤ 0.0001. Two-way ANOVA was performed.

**Figure EV4.**
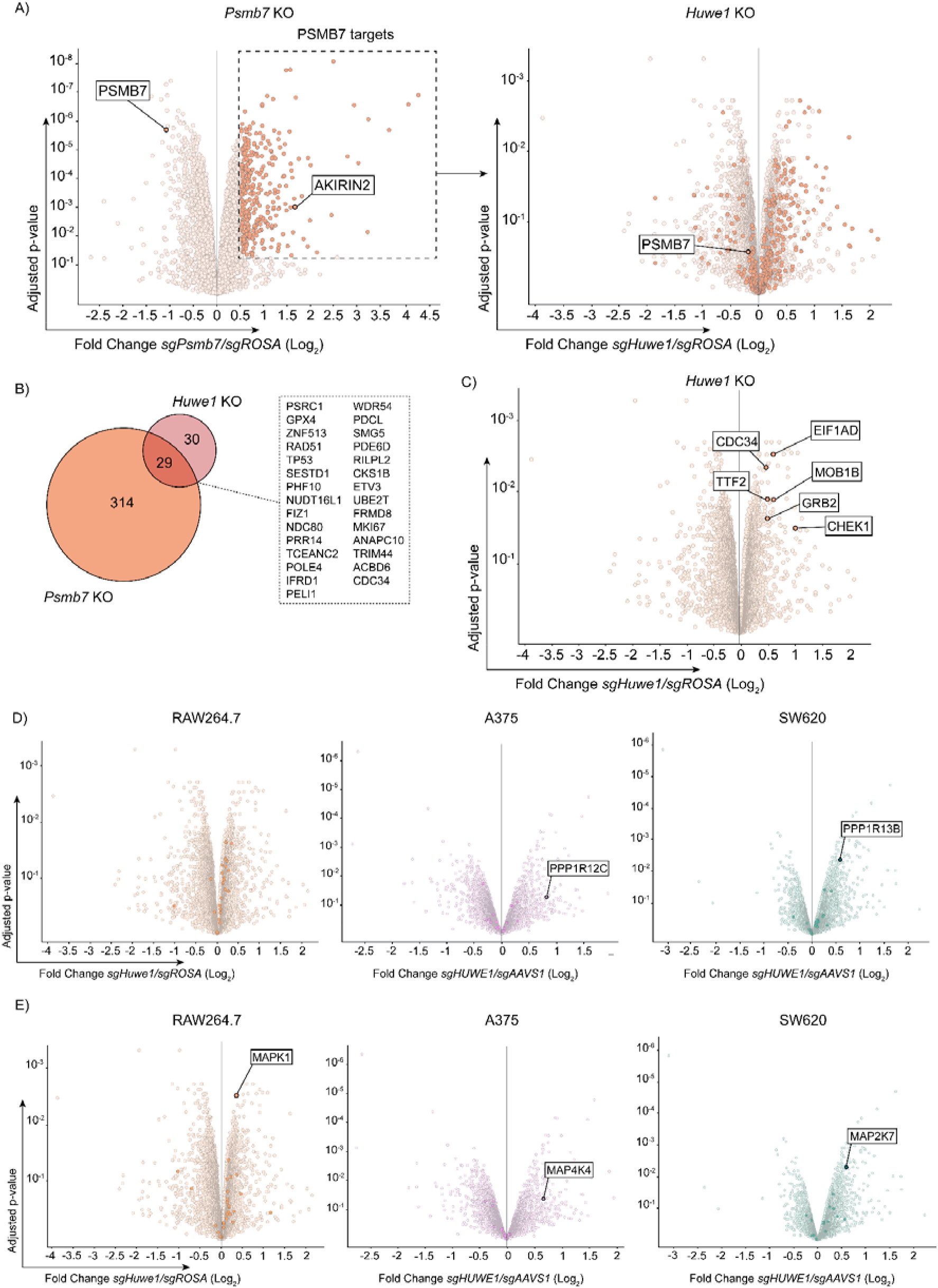
HUWE1 regulates TTP phosphorylation status, and thereby its TTP stability. **A-D)** RAW264.7-Dox-Cas9 cells expressing sg*ROSA* or sg*Huwe1* were treated with Dox for 5 days to induce Cas9. Cells were incubated with LPS for the indicated time points (min). Phosphorylation of **A)** p38, **B)** MK2, **C)** ERK, and **D)** JNK was determined by western blot. Total and phosphorylated kinase levels were quantified from non-saturated western blot signals and normalized to ACTIN. The ratio of the phosphorylated protein signals to their respective total protein signals was plotted. Data represent the mean and s.d.; n = 3. Two-way ANOVA was performed. **E)** RAW264.7-Dox-Cas9 cells expressing sg*ROSA* or sg*Huwe1* were treated with Dox for 5 days to induce Cas9. Cells were incubated with LPS for the indicated time points (h). Phosphorylation of p53 was assessed by western blot. As a control for the induction of p-p53, RAW264.7-Dox-Cas9 were treated with DNA damage inducer Etoposide for 6 h.

**Figure EV5.**
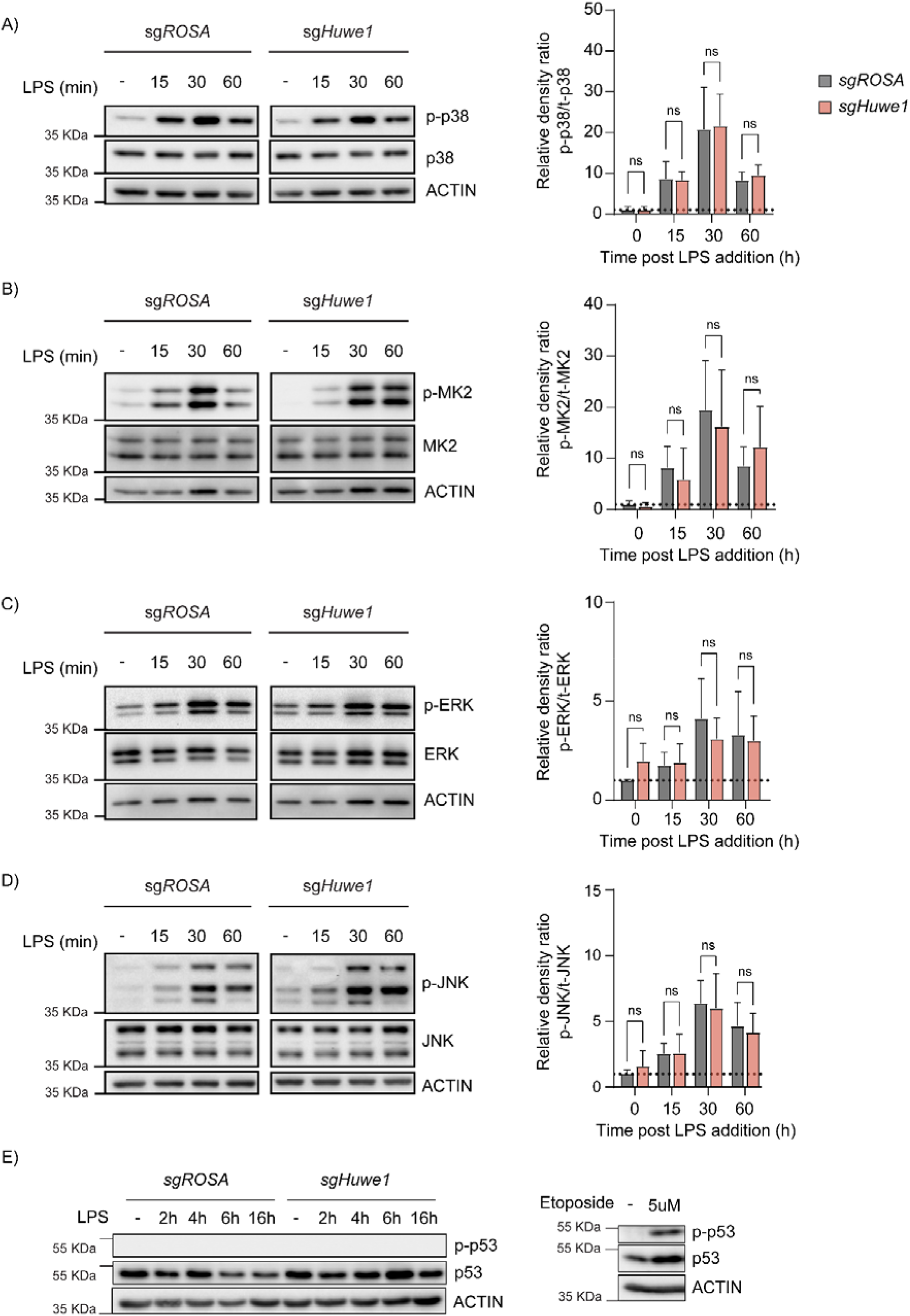
**A)** Proteome changes in RAW264.7-Dox-Cas9 cells expressing sg*Psmb7* by quantitative mass-spectrometry. Proteins classified as PSMB7 targets are highlighted in orange and displayed in the *Huwe1* knock-out volcano plot. (adjusted p-value ≤ 0.05 and Fold Change (Log_2_) ≥ 0.5; n = 3) **B)** Venn diagram showing the overlap in proteome changes of *Psmb7*- and *Huwe1*-targeted RAW264.7-Dox-Cas9 cells. Shared targets are listed. Twenty-nine proteins were identified as common targets (adjusted p-value ≤ 0.05 and Fold Change (Log_2_) ≥ 0.5; n = 3). **C)** Volcano plot representing proteome changes of *Huwe1*- and *ROSA*-targeted RAW264.7-Dox-Cas9 cells. Known HUWE1 targets are highlighted in orange (adjusted p-value ≤ 0.05 and Fold Change (Log_2_) ≥ 0.5; n = 3). **D-E)** Proteome changes of *Huwe1*-targeted RAW264.7, A375, and SW620 cell lines are plotted. PP1/2 proteins components in **D)**, and MAPK proteins in **E)**, are highlighted. Only significantly enriched proteins are labelled (adjusted p-value ≤ 0.05 and Fold Change (Log_2_) ≥ 0.5; n = 3).

**Figure EV6.**
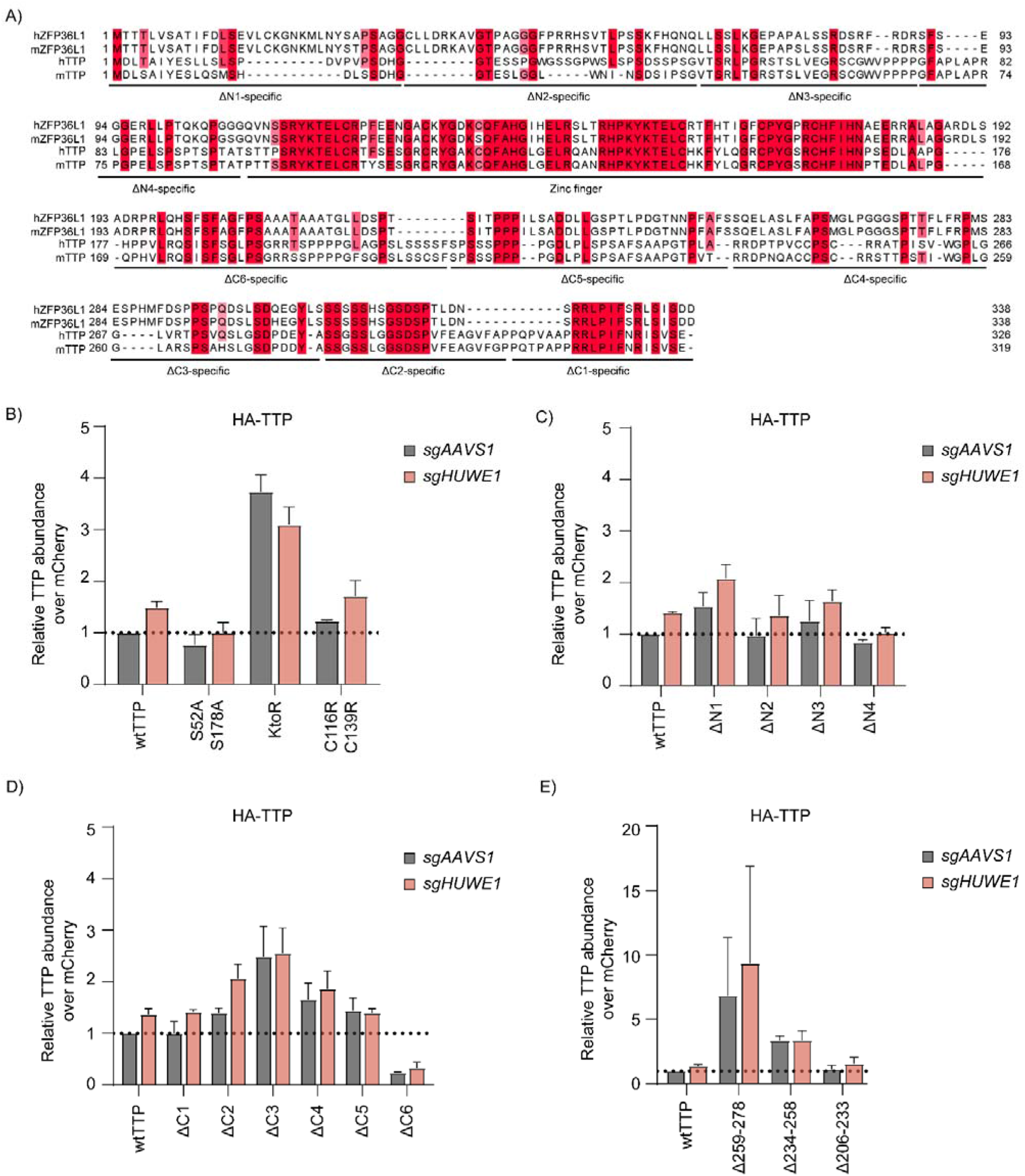
**A)** Alignment of mouse and human TTP and ZFP36L1 orthologs. **B)-E)** TTP protein levels from Fig.6 B)-E) were quantified from non-saturated western blot signals, normalized to the internal control mCherry, and plotted.

## Materials and methods

### Vectors

The lentiviral mouse genome-wide sgRNA library (six sgRNAs/gene) has been described previously (Michlits *et al*, 2020). Lentiviral vectors driving the expression of a single sgRNA or a dual sgRNA from a U6 promoter, and either eBFP2 or iRFP from a PGK promoter have been described previously (de Almeida *et al*, 2021). Single sgRNA CDSs were cloned in pLentiCRISPRv2 (Addgene plasmid 52961) to perform stable knock-outs in HEK293T cells. A Dox-inducible Cas9 lentiviral vector was modified from LT3GEPIR (Addgene plasmid 111177): T3G-GFP-(miR-E)-PGK-Puro-IRES-rtTA3, in which the GFP-mirE cassette was replaced by Cas9-P2A-GFP from pLentiCRISPRv2. The TTP stability reporter (pLX-SFFV-mCherry-TTP-P2A-BFP) was constructed by cloning the open reading frame (ORF) of murine TTP into a modified pLX303 vector (Addgene plasmid 25897). Lentiviral N-terminally HA-tagged-TTP deletions or point mutant variants were obtained by cloning the indicated variants of murine TTP ORF into a modified pLX303 vector. For this purpose, cDNAs encoding mTTP mutants were purchased from Twist Bioscience. All 3xHA-TTP constructs co-expressed mCherry through a P2A site to monitor protein expression and protein stability. All plasmids and sgRNAs used in this study are listed in the Appendix.

### Cell culture

All cell lines used in this study and their applications are listed in the Appendix. RAW264.7 Dox-Cas9 cells were generated by transducing RAW264.7 cells with pRRL-TRE3G-Cas9-P2A-GFP-PGK-IRES-rtTA3 lentiviral vector. Cas9 expression was induced with 500 ng/ml of Docycycline hyclate (Dox, Sigma-Aldrich, D9891) and single cells were sorted by FACS into 96-well plates using a FACSAria III cell sorter (BD Biosciences) to obtain single-cell-derived clones. Cas9 function and leakiness of the TRE3G promoter in the absence of Dox was tested in competitive proliferation assays. For mCherry-TTP reporter cells, pLX303-SFFV-mCherry-TTP-P2A-BFP was transduced into RAW264.7-Dox-Cas9 cells, and cells co-expressing mCherry and BFP were sorted by FACSAria III cell sorter into 96-well plates to obtain single-cell-derived clones. To obtain 3xHA-TTP expressing cells, pLX303-SFFV-mCherry-P2A-3xHA-TTP was transduced into RAW264.7-Dox-Cas9 cells, and cells expressing mCherry were bulk sorted using a FACSAria III. Bone marrow-derived macrophages (BMDM) were differentiated from bone marrow isolated from femurs and tibias of 8-to-12-week-old mice from Cas9 knock-in mice of both sexes (Platt *et al*, 2014). Femur and tibia marrow was centrifuged and cells were resuspended in DMEM. Cells were differentiated in DMEM (Sigma-Aldrich, D6429) containing recombinant M-CSF for 10 days. All cells were cultured at 37 °C and 5% CO_2_ in a humidified incubator. All animals were maintained in the pathogen-free animal facility of the Research Institute of Molecular Pathology, and all procedures were carried out according to an ethical animal license that is approved and regularly controlled by the Austrian Veterinary Authorities (License Number: GZ: 516079/2017/14).

### Transfections

All transfections were perfomed by mixing DNA and polyethylenimine (PEI, Polysciences, 23966) in a 1:3 ratio (μg DNA/μg PEI) in DMEM (Sigma-Aldrich, D6429) without supplements. Transfection was performed using 500 ng of total DNA. The day before transfection, 2×10^5^ HEK293T cells were seeded in 6-well clusters in fully supplemented media. Cells were harvested 48 hours after transfection, washed with ice cold PBS and and stored at −80 ºC until further processing.

### FACS-based CRISPR–Cas9 screens

The genome-wide Vienna sgRNA library was was lentivirally packaged in semiconfluent Lenti-X cells (Takara) via PEI transfection. Following double harvest, the collected supernatant was cleared of cellular debris by filtration through a 0.45 μm PES filter and stored a +4 ºC. The obtained virus was used to transduce RAW264.7 Dox-Cas9 cells at a multiplicity of infection (MOI) of less than 0.2 TU/cell, and 600 to 1,000-fold library representation. The percentage of library-positive cells was determined after 4 days of transduction by immunostaining of the Thy1.1 surface marker, and subsequent flow cytometric analysis. Library-positive cells were selected with G418 (1 mg/ml, Sigma-Aldrich, A1720) and expanded. Genome editing was induced with Dox (500 ng/ml, Sigma-Aldrich, D9891) and Cas9-GFP expression was monitored by FACS. Prior to Cas9 induction with Dox (Day 0), as well as before each FACS sort, an unsorted reference sample was collected. For this, a number of cells corresponding to at least 1,000-fold library representation was collected and stored at −80 ºC until further processing. After 3 days and 6 days of Cas9 induction, cells were sorted at FACS. Cells were harvested, washed with PBS and stained with Fixable Viability Dye eFluor (1:1,000, eBioscience, 65-0865-14) for 30 min. Subsequently, cells were washed three times with PBS, strained through a 35 µm nylon mesh and sorted in DMEM using the FACSAria II or FACSAria III cell sorters operated by BD FACSDiva software (v8.0). For the sort the following gating strategy was used: debris, doublets, dead (Viability Dye positive), Cas9-negative (GFP), mCherry- and BFP-negative cells were excluded. 5% of cells with the lowest and 1% of cells with the highest mCherry-TTP signal were sorted into PBS; same for the BFP internal control. At least 3 × 10^6^ (mCherry^low^ and BFP^low^) and 5 × 10^5^ (mCherry^high^ and BFP^high^) cells were collected for each time point. Sorted samples were re-analysed for purity, pelleted and stored at −80 ºC until further processing. The gating strategy for flow cytometric cell sorting is shown in Appendix Figure S2.

### Protein half-life determination

To estimate HA-TTP protein half-life, RAW264.7 Dox-Cas9 cells expressing sg*Huwe1* or sg*ROSA* were treated with Dox for 5 days before translational elongation was inhibited using 40 μg/ml of cycloheximide (CHX, Sigma-Aldrich, C1988). At indicated time points, whole cell lysates were prepared, analysed by western blot, quantified, and normalized to ACTIN levels and to time point 0 as indicated. Single exponential decay curves were determined using GraphPad Prism (v9), from which protein half-lives were calculated.

